# Implicitly Learned Higher Order Associations Differentiates Recent and Remote Retrieval of Temporal Order Memory

**DOI:** 10.1101/2021.07.03.451031

**Authors:** Shruti Shridhar, Vikram Pal Singh, Richa Bhatt, Sankhanava Kundu, J. Balaji

**Affiliations:** Center for Neurosciences, Indian Institute of Science, Bangalore – 560012

## Abstract

Memory of an ordered sequence of distinct events requires encoding the temporal order as well as the intervals that separates these events. In this study, using order place association task where the animal learns to associate the location of the food pellet to the order of entry into the event arena, we probe the nature of temporal order memory in mice. In our task, individual trials, become distinct events, as the animal is trained to form unique association between entry order and a correct location. The inter-trial intervals (> 30 mins) are chosen deliberately to minimise the working memory contributions. We develop this paradigm initially using 4 order place associates and later extend it to 5 paired associates. Our results show that animals not only acquire these explicit (entry order to place) associations but also higher order associations that can only be inferred implicitly from the temporal order of these events. As an indicator of such higher order learning during the probe trail the mice exhibit predominantly prospective errors that declines proportionally with temporal distance. On the other hand, prior to acquiring the sequence the retrospective errors are dominant. Additionally, we also tested the nature of such acquisitions when temporal order CS is presented along with flavour as a compound stimulus comprising of order and flavour both simultaneously being paired with location. Results from these experiments indicate that the animal learns both order-place and flavour-place associations. Comparing with pure order place training, we find that the additional flavour in compound training did not interfere with the ability of the animals to acquire the order place associations. When tested remotely, pure order place associations could be retrieved only after a reminder training. Further higher order associations representing the temporal relationship between the events is markedly absent in the remote time.

## Introduction

Lifetime experiences are characterised by events and are encoded in mammalian brain^1^. These events are then thought to be retrieved as episodes and hence the corresponding memories are termed as episodic memories^2–5^. This class of memory is thought to encode the contextual^6^ as well as temporal information^7,8^ content of the animal’s lifetime experiences. The temporal content constitutes both the information specific to the episode and the information that positions the episode in a sequence of events^9,10^. Temporal content can be characterised by measuring the independent parameters such as event boundary^11^, temporal order, duration of and the interval between these events^12–14^. Additionally, recency defined as the measure of time elapsed since the start of an event has been proposed and used to place the event in an allocentric frame of reference^15,16^. Thus, temporal memory and its characterisation is one of the intensely studied areas in the field of learning and memory. Temporal memory has been probed through different measures and attempts have been made to elucidate the behavioral correlates and their manifestations using model systems. Positioning the events with reference to external laboratory reference requires representation of time internally in an animal’s brain. Time cells is thought to provide one such mechanism. Studies involving time cells show that neural correlates of time in the Hippocampus, prefrontal cortex and basal ganglia^7,17–19^.

Interestingly, studies to probe the ability of the animals to encode an event sequence show that rodents can encode relative position of events in time. Such studies invariably involve a modified Go- NoGo task presenting the animal with two choices. Specifically, studies have been performed to investigate the memory for sequence of cue presentation inside of a trial^20–24^. Fortin et al. (2002) and Kesner et al. (2002) presented rats with unique sequences of odours and trained to select the odour which occurred earlier in the sequence. In these tasks the animal is exposed to a set of odours one followed by the other and later when presented with two choices, the rats are rewarded for choosing the one that occurred earlier in the sequence. Thus, demonstrating that the animals can encode the relative position of one odour with respect to other. Through such studies it has been established that hippocampus is required for identifying which of the odours comes earlier in a sequence. Recently in rodents schema formation for a set of sequences has been studied using modified Go-No Go task and the neural correlates representing the inter-relationship is shown to exist in OFC^25^. Again, the time interval between odour presentations in this study is short (4-8s).

However, temporal order studies utilising odours/events that are separated by 12 – 150 s have shown that animals can infer and retrieve the temporal distances when trained and probed using a 2-choice task. The training sessions in some of these tasks reward the choice of the odour/event that is presented earlier in the sequence. However, in such studies, the animals have to select from a choice of two odours/events without having to compare all the members of the sequence simultaneously. Such paradigms have also probed for temporal transitive inference, where rats are trained in pair-wise sequence associations (A>B, B>C, C>D, D>E) were tested for the formation of sequence relationships across these pairs (Specifically that of B>D) ^26^. While these studies have established that ordinal relationship in a sequence can be probed using a two-choice test, it is not clear if the animals can acquire all the elements of the sequence and their inter-relationships.

Using a modified object recognition task, Barker et al^27^ showed that the familiarity of object diminishes with time suggesting the representational memory of these objects decrease with time. Thus, they argue and show the familiarity can serve as a measure of elapsed time and hence can be used to encode order. Further this study has spaced apart presentations ranging from 5 minutes to 180 minutes within a sequence and have shown robust order memory. However, all the above paradigms be it a test using 2 forced choice probes or the test using Go – NoGo paradigms exhibit certain universal characteristics i) It is far more difficult for the subjects to discriminate between items adjacent to each other in the sequence ii) the rodents are capable of transitive inference when trained with pair-wise temporal sequences. These experiments entail learning through discrimination or disambiguation^23^ and differ greatly from free-recall paradigms to study sequence recall, which until recently have primarily been performed in human subjects. Fewer such studies exist in rodents, where Ishino et al ^28^ used a modified SRT task in rats to probe for recall of a 3-nose-poke sequence (from an option of 5 nose-poke holes, out of which they were trained only in 3) but the time scales involved are in the range of s well within the duration of the task and within the realms of working memory.

In humans, sequence memory has been studied and characterised extensively. It has been shown that when asked to enumerate the contents of an event, such as a list of words there is a strong recency effect. This results in the words that are being presented towards the end of the list being more likely to be remembered than other parts of the list. However most of these characterisation and studies are based on free recall or reaction times to wrong sequence presentation where we learn about the effects of temporal contiguity^29,30^.

In such tasks the events are separated by small time scales (∼ms) typically corresponding to working memory regimes. Thus, it is unclear how and if the temporal order of events separated by intervals corresponding to the short-term memory are encoded. A majority of the paradigms used to test temporal order memory do so, with smaller time delays between the presented objects/events ranging from few ms to minutes.

On the other hand, episodic memories which were once dependant on hippocampus are thought to be system consolidated^31–33^ wherein, they acquire the ability to be retrieved independent of hippocampus over time. In rodents through several behavioural paradigms the time scale of systems consolidation has been established to be ∼ 30 days^6^. During this process, memories tend to lose richness of detail, and generalize the stimuli and contextual information^34–36^. It has also been shown that it is not that the consolidated memory is unable to encode or retrieve the detailed information, but the loss is incidental^37^ and can be cue dependent^38^. It is interesting to note that temporal information needs to be acquired during encoding as recent studies have shown that acquisition of two memories close in time tend to encode them in overlapping population^39^ of neurons while temporally close retrieval of memories of two events that are separated in time does not result in such overlap^40^. Thus, it would of interest to know if the temporal order memory once formed can be retrieved remotely and if retrieved, is it different from the recent memory.

Given that much of our understanding of the sequence representation in the brain comes from studies that utilise limited number of choices little is known on the nature and the ability of the animal to encode the temporal order in a sequence. We wanted to ask if the animal is able to remember the absolute position of an event in a sequence when cued for ordinal position in a sequence. In this context in another paradigm after about 22 multiple training sessions, rats learn 6 flavour-place associations in an event-arena, and the memory can be retrieved even after the hippocampus has been lesioned while initially being dependant on it for its formation ^41^. We utilize this paradigm, for which the time-scale of consolidation has been established and investigate the nature of temporal memory resulting from sequence of distinct events separated over ∼45 mins (i.e outside of working memory regime).

Thus, in this study, we develop and use a flavour place association task to probe temporal order memory when a sequence of events is presented to the mice. Further we probe, if the mice can acquire not only this sequence but if the mice can also acquire the temporal interrelationship that exist between these events and if they are preserved during remote retrieval.

## Results

### Establishing the Event Arena based Order-Place Association paradigm in mice

We design and establish a novel task, based on event arena to probe temporal order memory in mice. Our behavioral paradigm for studying temporal order of events consists of 4-member order place associations (OPAs) and later we extend it to 5 OPAs. Detailed description of the setup and the behavioral design is presented in the methods section. Briefly, in our study the arena consists of a sand-filled square base, with a raised wall on the perimeter (Fig. 1A). Below the square base are 7 x7 grid of circular holes, of which any 4/5 positions are chosen for the task. This is done such that there is no rotational symmetry (please see S.Fig 1) in the placement of these holes. Once chosen these sand-wells are marked with a circular annulus, seen as white rings in Fig. 1A. These rings along with two distinct intra-maze cues (marked as visual cues1 and 2 in Fig. 1A) constitutes the ‘map’ of the arena. Once identified the holes are then fitted with sand-filled cups, at the base of which a food pellet is buried (Fig. 1A inset). 4 start boxes present along the edges, mark the 4 possible entry points into the arena. During each training session the mice are placed in the arena 4/5 times, and the order of entry (O_n_) determines the specific location (L_n_) at which the food pellet is buried. The food location is changed to one of the remaining non-baited well for subsequent entries in that session. These locations and entry order is kept consistent for a given mice all through the training thus forming a ‘O_n_ - L_n_’ pairing.

**Figure 1:**
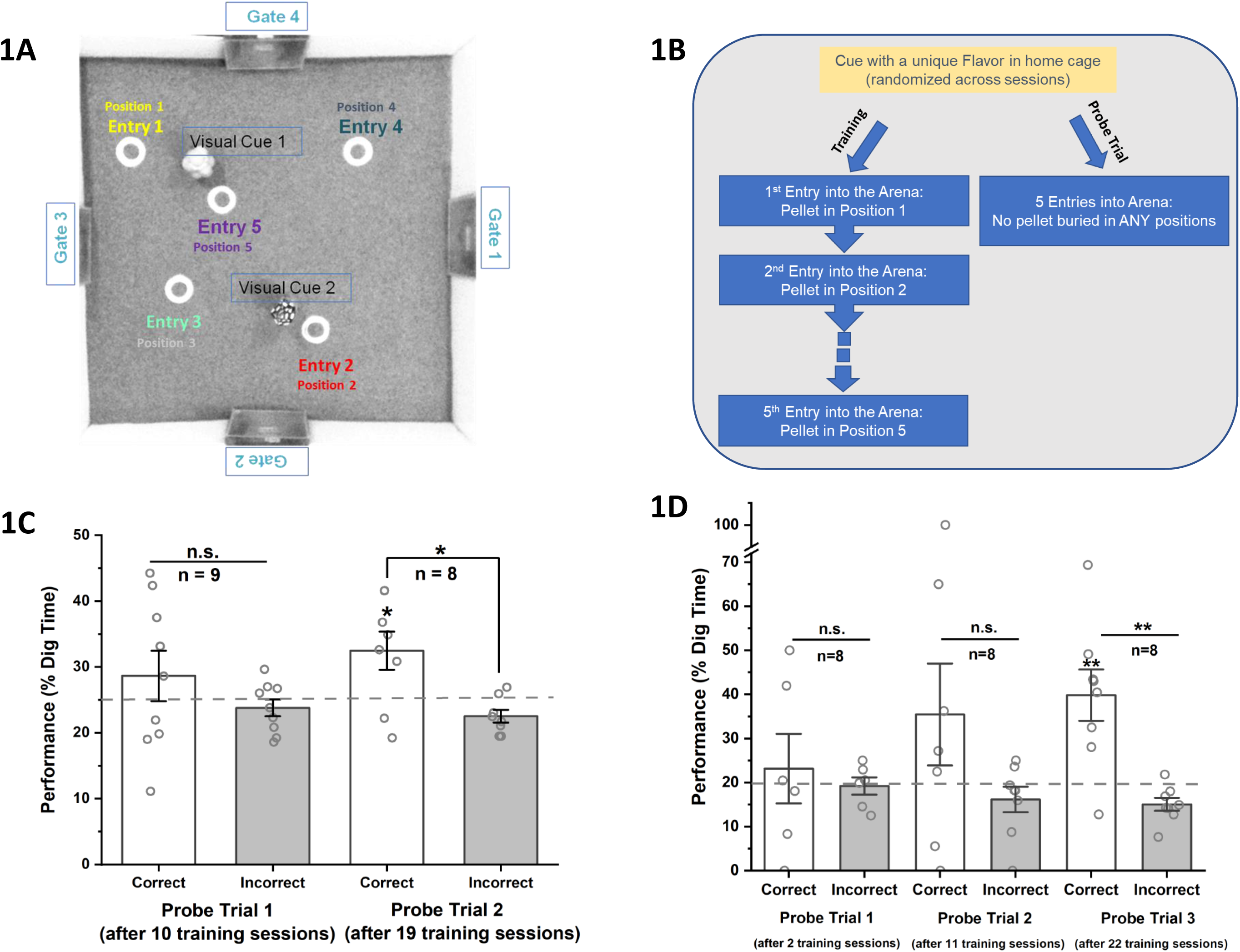
Event Arena based behavioral paradigm for temporal order memory (Order-Place Association, OPA Paradigm). **1A)** Birds-eye view of the event arena (constructed in-house) with 4 entry points (Gate 1-4), 5 sand-wells, seen as white annular rings and 2 visual cues. One of the 5 sand wells has an accessible flavor pellet in a given trial. **1B)** Schematic/flow chart of the behavioral training and probe trials; The behavioral training consists of multiple sessions across days, with each mice undergoing 5 training trials in a session. The location of each of the food pellet in each of these trials are different and are contingent across sessions for a given trial number. **1C)** Acquisition of temporal order memory following 4 Order-place association training probed during PT1 and PT2 conducted after 10 and 19 training sessions, respectively; Open circles are the mean % dig-time for a given mouse in the correct (open/white bars) and incorrect (grey bars) locations. The mean % dig-time for the incorrect and correct locations are not significantly different at PT1 (T=0.95 p=0.18 dF=8 N=9, paired t-test) but they are significantly different at PT2 (T=2.56 p<0.02 dF=7 N=8, paired t-test). The dotted grey line signifies the chance level of 25% for a 4-position task, and at PT2 the mice dig at the correct location significantly higher than chance (T= 2.56 p<0.02 dF =7 N=8, 1 sample t-test). **1D)** Acquisition of temporal order memory following a 5 Order-place association training probed during PT1, PT2 and PT3 conducted after 2,11 and 22 training sessions respectively; Open circles are the mean % dig-time for a given mouse in the correct (open/white bars) and incorrect (grey bars) locations. The mean % dig-time for the incorrect and correct locations are not significantly different at PT1 (T=0.39 p=0.71 dF=5 N=6, paired t-test) or at PT2 (T=1.33 p=0.11 dF=7 N=8, paired t-test) but they are significantly different at PT3 (T=3.39 p<0.006 dF=7 N=8, paired t-test). The dotted grey line signifies the chance level of 20% for a 5-position task, and at PT3 the mice dig at the correct location significantly higher than chance (T=3.39 p<0.006 dF=7 N=8, 1 sample t-test). The degree of significance of the means comparison are indicated as * with each * corresponding to an order of magnitude. * p<0.05 ** p<0.01 *** p<0.001 ****p<0.0001

The mice are initially habituated to the event arena and or food restricted to facilitate their digging to obtain food reward. Prior to training, the mice are shaped to dig for the food pellets and retrieve them from the bottom of the sand-well. Following which each training trial consisted of cueing the animal with a flavoured food pellet in their home cage for 30 seconds and then placing them in the arena. The order of the presented flavour is randomized. However, in a given session, 4/5 distinct flavours are used to facilitate the discrimination of different entry events in a given session. The choice of start boxes/entry point used is also randomized. In any given trial, the pellet would be buried only in one specific location corresponding to that entry order (O_n_), thus enabling the acquisition of association between the position of the baited sand well and the temporal order of entry. This pairing is maintained across the days as multiple training sessions are required to form the OPA.

The trials begin by placing the animal in the event arena after introducing the food pellet in the home cage. Once they are placed in the arena, the mice are permitted to explore and dig for 120 seconds. If they do not retrieve the food pellet within this time frame, they are shown the correct location at the end of the trial by the experimenter. The event arena is also sprayed with 70% EtOH, as we find the smell of alcohol masks the odour cues of the buried pellets and facilitates foraging based on spatial cues. The mice, during the initial training sessions dig in all locations but eventually learn to perform focused search and exhibit targeted digging.

The strength of the memory is measured by carrying out probe trials (PTs). These are conducted similar to the training session with crucial differences: 1.) The buried food pellet at the bottom of the sand well is not accessible to the animal, 2.) We leave the mice in the arena for 60 seconds (as against 120 seconds during training trials) and monitor its digging pattern. Extinction due to non-availability of food pellet during probe trails is avoided through presentation of food pellet in the correct location by the experimenter at the end of 60 s either through discretely burying the pellet at the correct location and leading the mouse to it or simply placing the pellet at the correct well.

Following 10 sessions of OPA training using 4 places, the mice are tested in the probe trail for memory (PT 1) the percentage time they spent digging in correct (29 ± 4 %) and incorrect location (24 ± 1.3%) are not significantly different (T= 0.95 p < 0.18 dF=8, N = 9, paired t-test). So, we trained the mice further. When tested after 9 more sessions of training, on the second probe trail (PT2) the mice spent significantly more time digging at the location of the correct well (32.45 ± 3 %) than the incorrect well (22.5 ± 0.97 %) (T =2.56 p < 0.02 dF=7, N = 8, paired t-test). We also see that the mice dig significantly above chance levels at PT2(T= 2.56 p < 0.02 dF =7, one sample t – test). These dig percentages are shown as grey (correct) and white (incorrect) bars in Fig. 1C where the open circles represent the average dig percent for each mouse at the correct and incorrect locations, while the average percent digging is represented by the bar. Taken together, we interpret this increase as arising from acquisition of stronger association between the order of entry and the food pellet location. The percentage digging time at the correct location is considered as a measure of their memory, with 25% dig time being the chance factor in a four-position task.

### Increasing the complexity of the Order-Place Association task

Given the ability of the mice to learn four OPAs we next investigate if mice would be able to acquire five OPAs. Despite a moderate increase of one extra position, such a pentameric set offered us possibility to probe intricate interdependences of the elements in a temporal order as described below. The mice now have 5 order-place associations to form, and the event arena map is structured as before, to avoid any rotational symmetry. We use 5 distinct flavors whose presentation is randomized, as is the entry point from session to session and the location of correct food well is contingent only upon the entry order (O_n_) as before. We interspersed 3 probe trials (PT1, PT2 and PT3) along the training sessions, and assessed the strength of the OPA memory. Our results from probe trails indicate that by PT3 (after a total of 22 training sessions) the animals have learned to associate the correct location of the food pellet based on the order of entry (T=3.39 p < 0.006 dF=7, N=8, paired t-test). These dig percentages are shown as grey (correct) and white (incorrect) bars in Fig. 1D where the open circles represent the average dig percent for each mouse at the correct and incorrect locations, while the average percent digging is represented by the bar. In PT3 the mice also dig significantly above the chance, which is 20% for a 5 position task (T=3.39 p < 0.006 dF=7, N=8, one sample t-test). Next, we proceed to test for signatures of temporal order memory from the errors the animals make. We reasoned that the nature of errors the animals make might be a good indicator for higher order associations that are acquired implicitly.

### Errors as measure of temporal order memory and notion of temporal rank in a sequence

The temporal order of events is characterised by the relative positions of the members in a sequence. Thus, given a sequence of ordered events {E_1_, E_2_, …, E_n_} present at ordinal positions {1^st^,2^nd^, n^th^} we define the absolute interposition difference(IpD_abs_) as |n - m|, IpD_abs_ measures the temporal distance or lag between events E_m_ and E_n_ and is purely a function of where the event occurs in a sequence and independent of spatial distance or absolute time of occurrence. Using such a measure we rank the positions (Fig 2A) such that Rank 1 corresponds to the correct location (IpD = 0) and Rank 2 corresponds to the location that immediately precedes or proceeds the correct location (IpD = 1) in the sequence and so on. In general, a location with rank n will be a correct location for an event that occurs either n-1 events after or before the current event. This enables us to classify the errors the mice make according to different ranks and ask if the propensity of making errors is different across the ranks.

**Figure 2:**
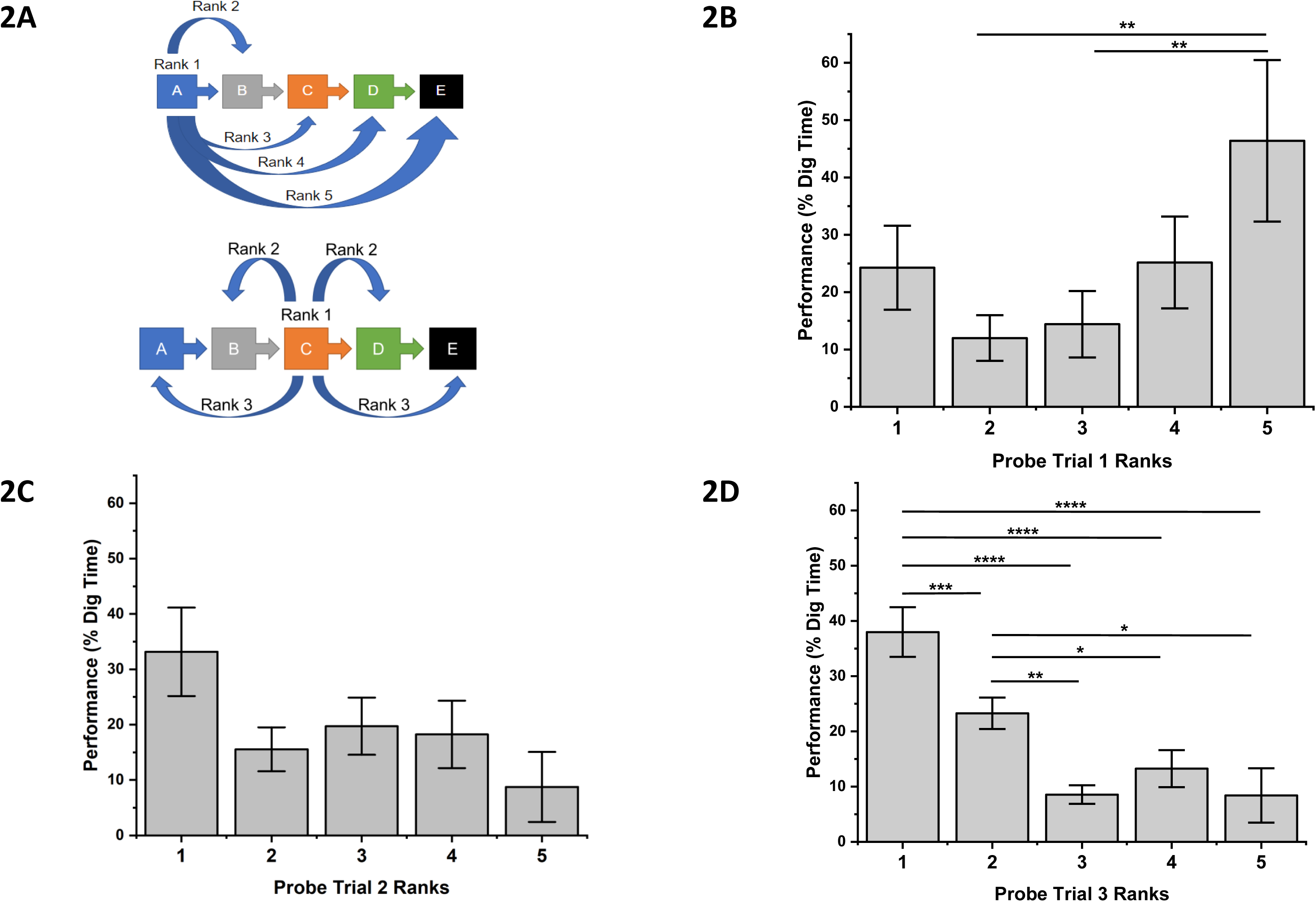
Measuring higher order associations in temporal order memory using errors. **2A)** Illustration of relationship among the events in a temporal sequence; Different colored squares, A to E, represent unique events presented in a sequence. Rank of an error made by the mouse is determined by the distance in the sequence between the location in which it has erroneously dug in and the correct location. We illustrate 2 sets of such ranks for the location corresponding to event A and C. Rank 1 corresponds to the correct location. **2B-D)**The grey bars in panels 2B,2C and 2D represent the mean % dig-time across ranks (1 to 5) measured during probe trials PT1, PT2 and PT3 in that order. The probe trials are conducted after 2, 11 and 22 training sessions. One-way ANNOVA performed at each of the probe trials revealed that the mean % Digging time across the ranks is different for PT1 (F= 2.7 p<0.032 dF=4, N=8) and PT3 (F=12.99 p<0.0001 dF=4, N=8) However, post-hoc analysis showed that only in PT3, Rank1 (the correct location) is consistently higher than all the other ranks (Fischer LSD post-hoc Rank1>2 p<0.0008, Rank1>3 p<0.0001, Rank1>4 p<0.0001, Rank1>5 p<0.0001) while Rank 2 digging is significantly higher than all the subsequent ranks (Fischer LSD Post-hoc Rank2>3 p<0.0004, Rank2>4 p<0.032, Rank2>5 p<0.02). In PT1 post-hoc tests reveal that it is Rank5 (incorrect locations which are the most temporally distant from the correct location) which is significantly greater than temporally closer Rank2 and Rank3 (Fischer LSD post-hoc Rank5>2 p<0.0031, Rank5>3 p<0.0072). The degree of significance of the means comparison are indicated as * with each * corresponding to an order of magnitude. * p<0.05 ** p<0.01 *** p<0.001 ****p<0.0001

Using the above ranking system, we segregated the digging shown by these animals at probe trials PT1 (Fig. 2B), PT2(Fig. 2C) and PT3 (Fig. 2D). ANOVA performed on the percent digging as function of ranks observed at PT3 showed a main effect (F = 12.99 p<0.0001) . Post hoc analysis showed a significantly increased digging at the Rank 1 position indicating the animal has acquired the memory for the correct location (Fisher LSD Post-hoc Rank1>2 p<0.0008, Rank1>3 p<0.0001, Rank1>4 p<0.0001, Rank1>5 p<0.0001). For detailed comparison refer to S.Table 1). In addition, the analysis also showed that Rank2 digging is significantly more than Rank 3 and the other subsequent ranks (Fisher LSD Post-hoc Rank2>3 p<0.0004, Rank2>4 p<0.032, Rank2>5 p<0.02). However, the difference between Ranks 3 and above are not significant. We see that the animals not only remember the correct location (Rank 1 digging), but the decline in percent digging is graded. We see the animals apart from digging at the correct location also dig at other locations and are more likely to dig at locations that are adjacent in sequence to the correct location, i.e IpD = 1 more than other locations with IpD >1 . We interpret this result as suggestive of the animals expressing temporal order memory to a first degree (Rank 2).

### The errors are further dissociated into forward and backward errors

Further to ranking the incorrect food locations in terms of their temporal distance defined as above we inquire if the errors reflect prospective and retrospective nature of search exhibited by the mice. Thus, we further classify the errors as “Backward/Negative” and “Forward/Positive” Errors. Backward errors measure the tendency of the mice to revisit previously rewarded locations while forward errors measure the tendency of mice to search in hitherto non-baited locations in that session. Thus, for instance if the mouse has entered for the third time in the training session for the day (Fig. 2A bottom), Position 3 corresponds to the correct sand-well location (orange square and marked as C) where the food reward is buried. It has already been trained for Position 1 and 2 (corresponding to the First and Second time it entered into the arena) and has not yet encountered Position 4 and 5. In such a scenario, digging in Position 1 and 2 constitute backward errors as they precede the correct location in the temporal sequence (n-m is negative). Digging in Position 4 and 5 constitute forward/positive errors as they proceed (and are to the right of) the correct location in the temporal sequence (n-m is positive). We observe that as the training progresses the mice make significantly more forward errors compared to backward errors and at PT3 we see forward errors are significantly more (T=3.93 p<0.0001 dF=122) (Fig 3H). The bar graph in Fig 3H shows the forward errors in dark grey and the backward errors in light grey across. Percent dig time from earlier probe trails show that this is not the case for PT1 (T=-0.84 p=0.8 dF=82), PT2 (T=-0.31 p=0.6 dF=106). While overall increase in the forward errors would be indicative of elimination strategy used by the animal to fetch the pellets, we see that the errors are higher for temporally close locations, again suggestive of animal implicitly acquiring the temporal order information.

**Figure 3:**
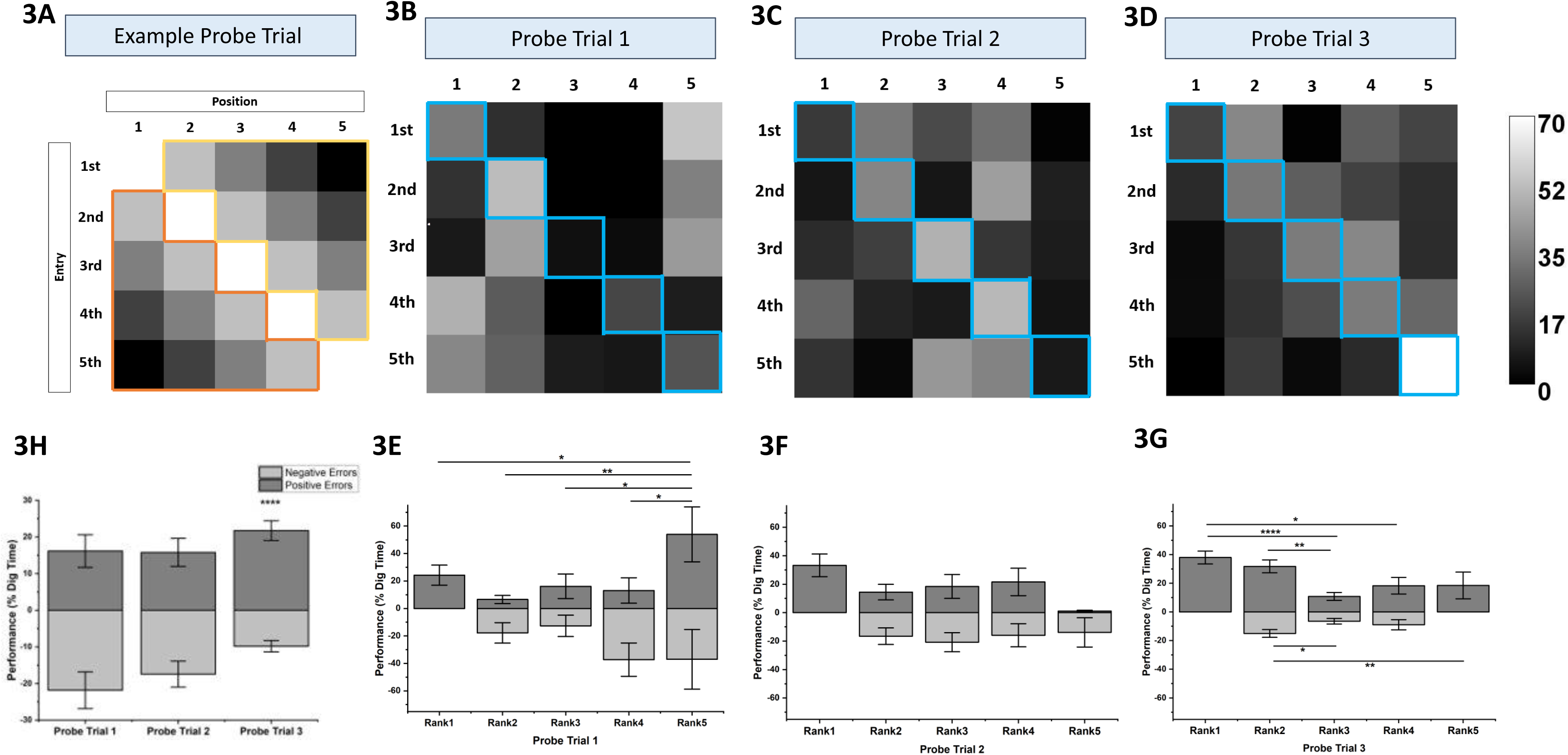
Measuring higher order associations in temporal order memory (OPA paradigm) using forward and backward errors. **3A)** Schematic illustration of a 5 × 5 matrix, where the rows represent the order of entry into the arena and the columns represent the locations/positions in the temporal sequence. The 5 diagonal elements represent the correct location, and every other location on either side of the diagonal (the remaining 20 elements/locations) corresponds to an error. The set of errors outlined in yellow correspond to forward/positive errors. The set of errors outlined in orange correspond to backward/negative errors, where the mouse digs in positions preceding those in the temporal order (and those which it has encountered before in the current training/probing session). This example figure shows an example heatmap, where the mouse has learnt the order-place associations. The diagonal shows the highest amount of digging. The positions adjacent to the correct location show reduced digging in a graded way, depending on the temporal distance from the correct location. In this example figure, the digging reduces in the incorrect locations, by the same measure, in the forward/positive and backward/negative directions. Intensity at each of the 25 elements represents the average performance (as percentage dig-time) for all mice at that position for a particular nth entry. No normalization has been performed. **3B-D)** The diagonal elements, outlined in blue, represent the correct locations. As the training progresses, the probes show higher digging in the correct locations. There is also a progressive increase in digging in the more forward/positive error locations compared to the backward/negative errors. The scale bar shows the lightest/highest % dig-time represented in the PT1(3B),PT2(3C),PT3(3D) heatmaps to be around 70% and the darkest/lowest % dig-time to be 0%. **3E-G)** The dark grey bars represent mean % dig time in locations representing Rank1 and rank-wise forward errors and the light grey bars represent rank-wise backward errors across PT1(3E), PT2(3F) and PT3(3G) respectively. One-way ANNOVA performed at each of the probe trials revealed that the mean % digging across the forward/positive ranks is different for PT1 (F=2.86 p<0.031 dF=4, N=8) and PT3 (F= 5.60 p<0.0005 dF=4, N=8) but not PT2 (F=1.48 p=0.22 dF=4, N-8). However post-hoc tests of the forward ranks of PT1 (3E) reveal that it is Rank5 (incorrect locations which are the most temporally distant from the correct location) which is significantly greater than the correct location, Rank1 and it’s temporally closer locations (Fischer LSD post-hoc Rank5+>1 p<0.05, Rank5+>2+ p<0.003, Rank5+>3+ p<0.02, Rank5+>4+ p<0.02). Post-hoc tests in PT3(3G) Forward/positive ranks reveal mean % dig time in Rank1(the correct location) is significantly higher in subsequent forward ranks but not in the immediately proceeding Rank2+ locations(Fischer LSD post-hoc Rank1>3+ p<0.0001, Rank1>4+ p<0.0021, Rank2+>3+ p<0.011). **3H)** The dark grey bars represent mean % dig time in locations representing total forward errors and the light grey bars represent total backward errors across PT1, PT2 and PT3. As the training progresses, PT3 shows forward errors are significantly higher than backward errors (T=3.93 p<0.0001 dF=122, N=8, t-test). The degree of significance of the means comparison are indicated as * with each * corresponding to an order of magnitude. * p<0.05 ** p<0.01 *** p<0.001 ****p<0.0001

When the percentage digging in forward and backward errors are further dissociated into ranks and analysed using ANOVA, only in PT3 (Fig 3G)do we see a main effect (F= 5.60 p<0.0005) where Rank 1 is higher than all subsequent forward ranks, except the immediately proceeding Rank 2+ (Fisher LSD post-hoc Rank1>3+ p<0.0001, Rank1>4+ p<0.0021, Rank2+>3+ p<0.011). ANOVA performed on the forward ranks in PT2 (Fig 3F) show no main effect, and in PT1 (Fig 3E) digging in Rank 5 +, which is incorrect (and the most distant in a temporal relationship from Rank1/Correct location) is higher than all other ranks(F=2.86 p<0.031 Fisher LSD post-hoc Rank5+>1 p<0.05, Rank5+>2+ p<0.003, Rank5+>3+ p<0.02, Rank5+>4+ p<0.02).

### A combined Flavor-Order Place Association task

Next, we inquire if such implicit acquisitions can be acquired even in the presence of distinct and explicit training. We do this through a simultaneous order-place and flavour-place association training. The mice are presented with 5 unique flavors in a fixed order, in every training session. While the start box entries are randomized as in the paradigm described before, the flavors are not randomized. The mice learned to associate BOTH the pellet flavour and the order/entry trial number with a specific location in the arena. We note that even though the order is maintained constant for a given mouse, they are different across the mice as described in methods. We design the test structure to probe different aspects of the dual association (Flavor and Order combined) memory that is being acquired. The probe trial proceeds by first cueing the mice with the flavour in the home cage and subsequently measuring the digging performance in the arena. The cueing and the tests are done by maintaining the same temporal order as that of training. Since in such probe trails both the flavour -place as well as order place associations are tested, we term them as whole probe trial (PTW). We also assessed for the temporal order memory alone. We do this through conducting explicit order-place association probes. In these Order probes we cue the mice with novel flavors (that are different from flavors used during training) and measure the time of digging at different positions. In these tests the mice need to depend only on the order of entry as a cue and such probe trials are termed as probe trails for order (PTO). Both PTW and PTO are conducted on consecutive sessions. After a total of 21 training sessions the mice dug in the correct location significantly higher than by chance(T=9.84 p<0.0001 dF=11 N=12, one-sample t-test) and significantly higher than the incorrect locations (T=9.84 p<0.0001 dF=11 N=12, paired t-test) as seen in Fig 4A. In the order probes conducted immediately after PT3W, in PT3O(Fig 4B), the mice showed higher digging in the correct locations than by chance (T= 4.41 p<0.0006 dF=11, N=12, one-sample t-test) and also higher than in incorrect locations (T=4.41 p<0.0006 dF=11 N=12, paired t-test), indicating that the mice are able to dig in the correct location even without the flavor cues.

**Figure 4:**
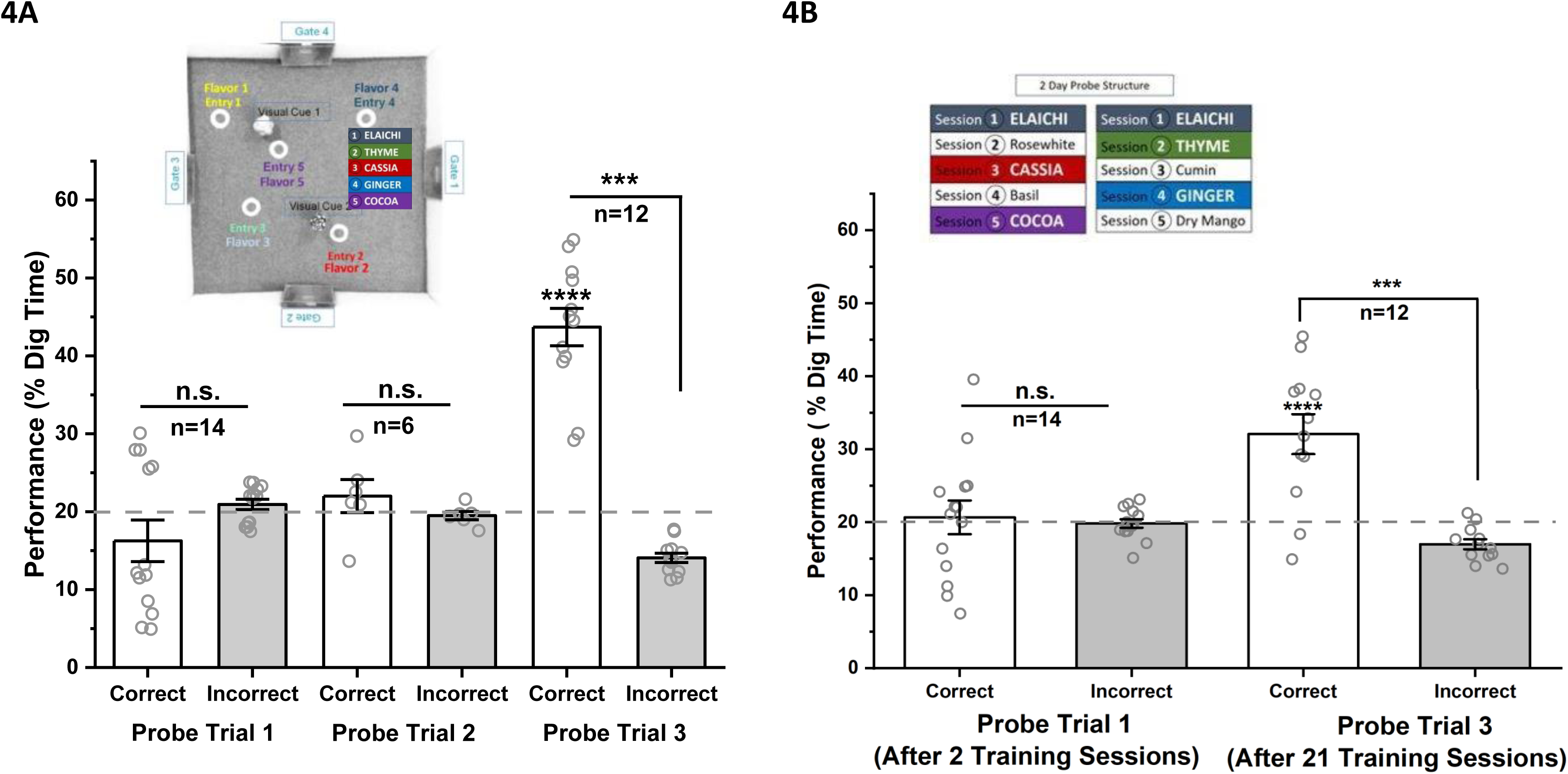
Combined Order-Flavor place association event arena task. **4A)** Inset shows a birds-eye view of the event arena with 5 sand-wells, seen as white annular rings and 2 visual cues. The 5 positions correspond to a particular entry-order and a unique flavor. Graph shows acquisition of temporal order memory following 5 Ordered-flavor-place association training probed during PT1, PT2 and PT3 conducted after 3, 15 and 21 training sessions respectively. Open circles are the mean % dig-time for a given mouse in the correct (open/white bars) and incorrect (grey bars) locations. The mean % dig-time for the incorrect and correct locations are not significantly different at PT1 (T= -1.40 p=0.91, n=14, paired t-test) and PT2 (T=0.942 p=0.19 dF=5 N=6, paired t-test) but they are significantly different at PT3 (T=9.84 p<0.0001 dF=11 N=12, paired t-test). The dotted grey line signifies the chance level of 20% for a 5-position task, and at PT3 the mice dig at the correct location significantly higher than chance (T=9.84 p<0.0001 dF=11 N=12, one-sample t-test). **4B**) Inset shows the structure of Order probes of the 5-flavor-order-place association paradigm. The probe is divided across two days, where trained flavor cues (Colored rectangles for the flavors Elaichi, Thyme, Cassia, Ginger, Cocoa) are alternatively replaced with novel flavors (white rectangles Rosewhite, Cumin, Basil, Dry Mango). Graph shows the acquisition of temporal order memory following training probed during PT1 and PT3 conducted after 3 and a total of 21 training sessions respectively. Open circles are the mean % dig-time for a given mouse in the correct (open/white bars) and incorrect (grey bars) locations. The mean % dig-time for the incorrect and correct locations are not significantly different at PT1 (T= 0.29 p=0.39 dF=13 N=14, paired t-test) but they are significantly different at PT3 (T=4.41 p<0.0006 dF=11 N=12, paired t-test). The dotted grey line signifies the chance level of 20% for a 5-position task, and at PT3 the mice dig at the correct location significantly higher than chance (T= 4.41 p<0.0001 dF=11 N=12, one-sample t-test). The degree of significance of the means comparison are indicated as * with each * corresponding to an order of magnitude. * p<0.05 ** p<0.01 *** p<0.001 ****p<0.0001

The percentage digging at different locations classified according to the temporal distance is presented in Fig. 5 as bar graphs for PT1W(5a), PT1O(5b), PT3W(5c) and PT3O(5d). The data for second probe trails PT2W are presented in SFig 2. The mice dug specifically in the correct location (Rank1) only during third probe trail. The ANOVA performed on the percent digging at each of the probe trial revealed a main effect in PT1O (F=3.42 p<0.01 dF=4, N=14) and in PT3 for both PT3W (F= 29.53 p<0.0001 dF=4, N=12) and PT3O(F= 9.05 p<0.0001 dF=4, N=12). PT1O shows higher digging in incorrect locations Rank3 and Rank5 (Fisher LSD post-hoc Rank3<2 p<0.01, Rank3>Rank4 p<0.003, Rank5>Rank4 p<0.02). The post hoc analysis for the final probe trials (PT3s) showed Rank1 percent digging is significantly higher than all other ranks for both PT3W (Fisher LSD post-hoc Rank 1>2 p<0.0001, Rank1>3 p<0.0001, Rank1>4 p<0.0001, Rank1>5 p<0.0001) and PT3O(Fisher LSD post-hoc Rank 1>2 p<0.0007, Rank1>3 p<0.0001, Rank1>3 p<0.0001, Rank1>4 p<0.0001 Rank1>5 p<0.003). Further post-hoc tests in the Order probe we see the rank 2 digging is significantly different from rank 3 (Rank2/3 p<0.017) as well as rank 4 (Rank2/4 p<0.03). However, this is not the case in PT3W, suggesting that the animals inferred and acquired the sequence information during these trials even in the presence of explicit training cues for flavour.

**Figure 5:**
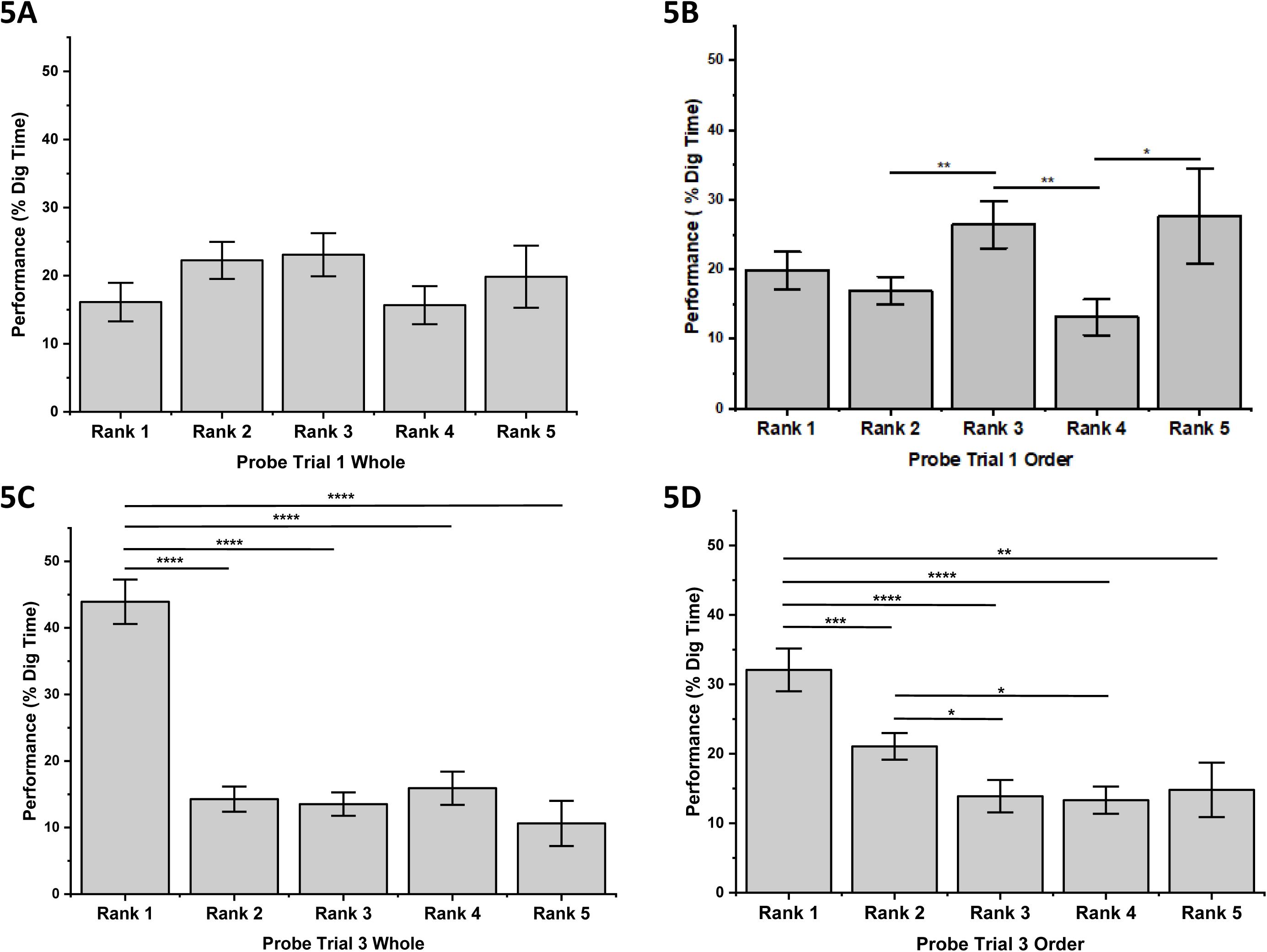
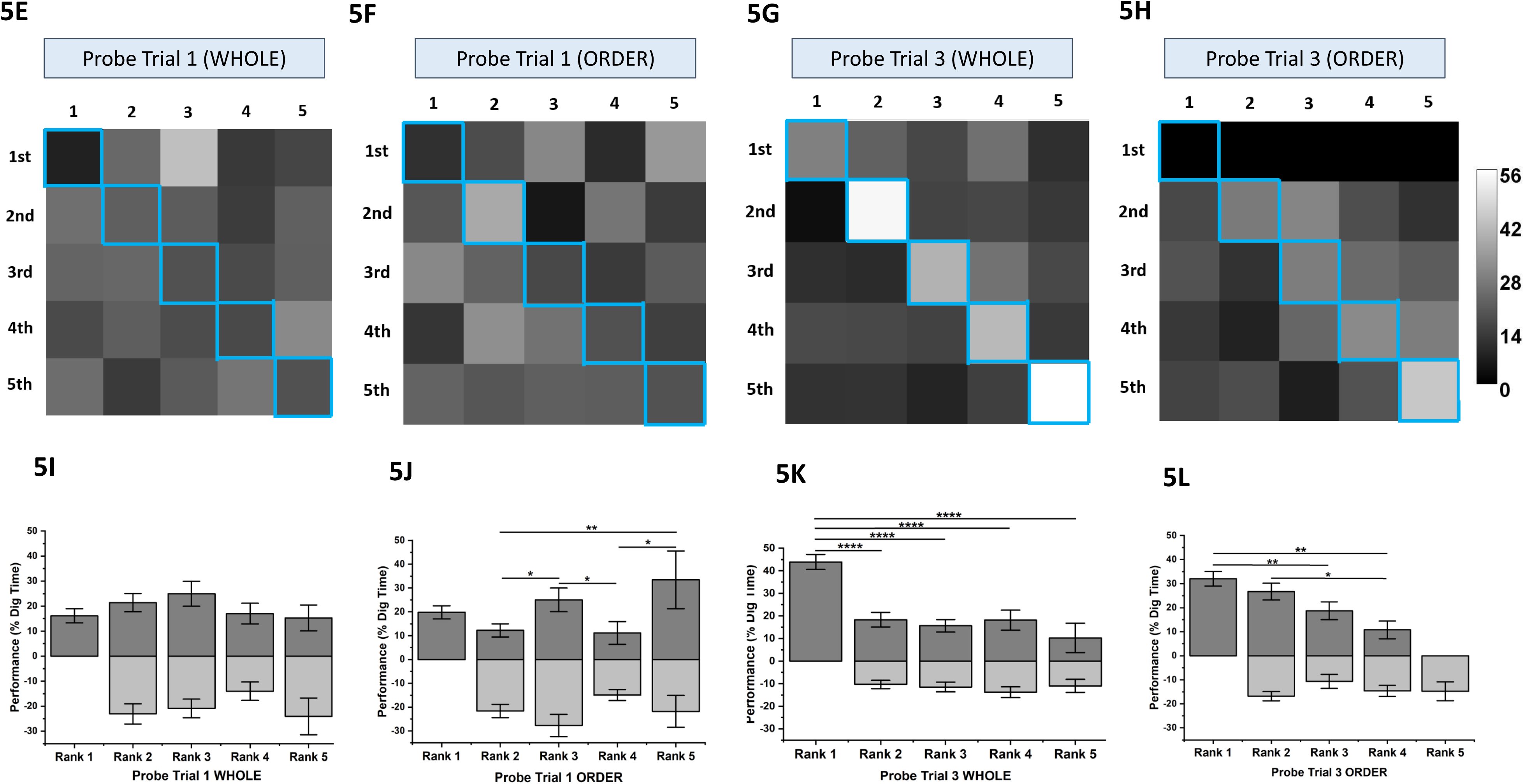
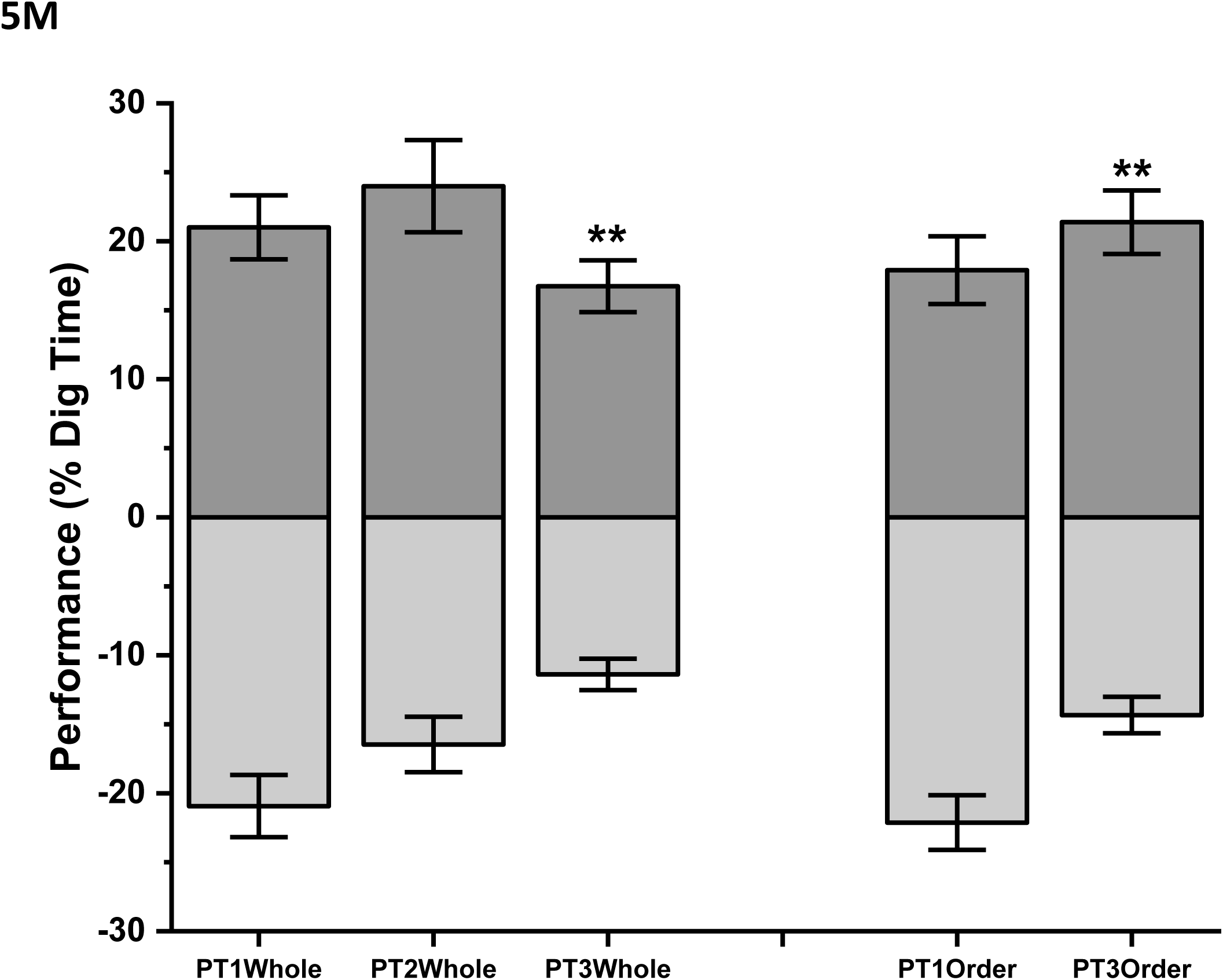
Measuring higher order associations in Order-Flavor Place Association (OFPA) task using errors. **Fig 5A-D**) The grey bars in panels 6A and 6B represent the mean % dig-time across ranks (1 to 5) measured during PT1 while cueing for flavor in order (Probe trial 1 Whole) and cueing with novel flavors (Probe trial 1 Order). The grey bars in panels 6C and 6D represent the mean % dig-time across ranks (1 to 5) measured during PT1 while cueing for flavor in order (Probe trial 3 Whole) and cueing without flavor (Probe trial 3 Order). PT1 and PT3 were conducted after 3 and 21 training sessions. One-way ANNOVA performed at each of the probe trials revealed that the mean % digging across the ranks is different for PT3 for whole probes (F= 29.53 p<0.0001 dF=4, N=12) and PT1O(F=3.42 p<0.01 dF=4, N=14) and PT3O (F= 9.05 p<0.0001 dF=4, N=12) for order probes. Post-hoc analysis for PT1O shows the incorrect ranks, Rank3 and Rank5 to be higher than others (Fischer LSD post-hoc Rank3<2 p<0.01, Rank3>Rank4 p<0.003, Rank5>Rank4 p<0.02). Post-hoc analysis for PT3 shows Rank1 to be significantly higher than all subsequent ranks for both PT3W (Fischer LSD post-hoc Rank 1>2 p<0.0001, Rank1>3 p<0.0001, Rank1>3 p<0.0001, Rank1>5 p<0.0001) and PT3O (Fischer LSD post-hoc Rank 1>2 p<0.0007, Rank1>3 p<0.0001, Rank1>3 p<0.0001, Rank1>4 p<0.0001, Rank1>5 p<0.003). Further post-hoc tests for PT3O show that Rank2 digging is significantly higher than Rank 3 (Rank2>3 p<0.017) as well as Rank 4 (Rank2>4 p<0.03). **5E-H)** Heatmaps of probes of OFPA paradigm;5E and 5F show heatmaps representing the % dig-time in the first probe trials, PT1W and PT1O respectively. 5G and 5H show heatmaps representing the % dig-time in the final probe trials, PT3W and PT3O respectively. The first probe trials show non-specific digging in all positions, the final probe trials show concentrated digging along the diagonal. Comparison of the final whole and order probes shows more concentrated digging along the diagonal in PT3W than in PT3O. The first row in PT3O, in 5H is left blank, because the mice are not probed for Position 1 corresponding to Elaichi/Cardamom and were cued with the flavor. The diagonal elements, outlined in blue, represent the correct locations. The scale bar shows the lightest/highest % dig-time represented in the PT1W, PT1O, PT3W, PT3O heatmaps to be around 56% and the darkest/lowest % dig-time to be 0%. **5I-J)** The dark grey bars represent mean % dig time in locations representing Rank1 and Rank-wise forward errors and the light grey bars represent Rank-wise backward errors across Probe trial 1 for Whole (PT1W) and Order(PT1O) conducted after 3 training sessions. One-way ANNOVA performed at each of the probe trials revealed that the mean % digging across the forward/positive ranks is not different for PT1W but it is for PT1O (F=3.09 p<0.02 dF=4, N=14). However post-hoc tests reveal than Rank1, the correct rank, is not higher than the subsequent forward ranks (Fischer LSD post-hoc Rank3+>2+ p<0.02, Rank3+>4+ p<0.035, Rank5+>Rank4+ p<0.02, Rank5+>Rank2+ p<0.009). **5K-L)** The dark grey bars represent mean % dig time in locations representing Rank1 and Rank-wise forward errors and the light grey bars represent Rank-wise backward errors across Probe trial 1 for Whole (PT3W) and Order(PT3O) conducted after a total of 21 training sessions. One-way ANNOVA performed at each of the probe trials revealed that the mean % digging across the forward/positive ranks is different for both PT3W and PT3O (F = 4.91 p<0.004 dF=3, N=12). In PT3W post-hoc tests reveal Rank1 to be significantly higher than all subsequent forward ranks (Fischer LSD post-hoc Rank1>2+ p<0.0001, Rank1>3+ p<0.0001, Rank1>4+ p<0.0001, Rank1>5+ p<0.0001). In PT3O post-hoc tests reveal that the mean % dig time with subsequent forward/positive ranks decreases (Fischer LSD post-hoc Rank1>3+ p<0.009, 1>4+ p<0.002 and 2+>4+ p<0.02). **5M)** The dark grey bars represent mean % dig time in locations representing total forward errors and the light grey bars represent total backward errors across PT1Whole, PT2whole and PT3whole and PT1Order and PT3Order. PT3 for both whole(T=2.46 P<0.008 dF=223 N=12, t-test) and order(T=2.86 p<0.003 dF=190, N=12, t-test) shows forward errors are significantly higher than backward errors. The degree of significance of the means comparison are indicated as * with each * corresponding to an order of magnitude. * p<0.05 ** p<0.01 *** p<0.001 ****p<0.0001

Further during our experiments, we notice that the animals progressively learn to eliminate the already baited locations. An indicator of this effect is the comparison of forward and backward errors. Fig. 5M shows these errors during different probe trails and we notice that both for whole as well as order probes the animals make more of forward errors compared to backward errors (PT3W T=2.46 P<0.008 dF=223, PT3O T=2.86 p<0.003 dF=190, t-test) only at probe trail 3 for both Whole and Order probes. However, when these errors are further distributed according to ranks we see (Fig. 5 I – 5L) a progressive decline in the digging percentage in the forward ranks of PT3O. ANOVA performed to compare the forward ranking errors shows the percentage digging across the ranks are significantly different in PT30(F = 4.91 p<0.004). Post-hoc analysis reveals that only Rank1-3+(Fisher LSD post-hoc p<0.009), 1-4+(Fisher LSD post-hoc p<0.002) and 2+-4+(Fisher LSD post-hoc p<0.02) are significantly different and not adjacent ranks (1-2, 2-3, 3-4), thus generating a staircasing effect wherein the reduction in percentage digging is more graded across ranks. Such a progressive decline is absent in the forward ranks of PT3W, where percentage digging in Rank1 is higher than all other ranks (F=14.87 p<0.0001, Fisher LSD post-hoc Rank1>2+ p<0.0001, Rank1>3+ p<0.0001, Rank1>4+ p<0.0001, Rank1>5+ p<0.0001). Rank2+,3+,4+,5+ showed no difference amongst each other in PT3W. We interpret these results as indicative of the animal exhibiting the acquired higher order associations amongst the locations when probed for order. We note that the animals do not exhibit such digging when probed for order along with the explicit flavour cue.

### Remote Order-Place association memory

Given that the training paradigm uses a sequence of events, and the animals acquire the temporal order present in that sequence through higher-order associations, we proceeded to ask if such information is preserved across time. We do this by probing the animal 30 days after training (remote time point). The white and the grey bars (Figure 6a) show the percentage digging exhibited by the animals in the correct and incorrect locations when tested (I) immediately after 21 training sessions (recent trial PT3) (ii) Tested 30 days after the final 23^th^ training session (Remote Probe Trial 1, RPT1) (iii) following a remote reminder training (Remote Probe Trial 2, RPT2) and (iv) 15 days following the remote reminder training(Remote Probe Trial 3, RPT3). When probed in RPT1 the animals do not exhibit (Fig 6a) correct location specific digging, which they did when tested after recent training. This could be due to the task being very strenuous and a gap of 30 days could lead to loss of familiarity with the arena and the task, leading to lack of performance rather than loss of order place association memory. We addressed this by performing a single training session (a reminder), and then conducting a probe trial. Re-training with a single session acts as a sufficient reminder and the % digging performance of the mice at the correct location is above chance (t= 3.72 p<0.005 dF=6 N=7, one-sample t-test)and higher than in the incorrect locations (t=3.72 p<0.005 dF=6 N=7, paired t-test) . Another probe trial conducted 15 days after that also shows significantly higher % digging performance in the correct locations than in the incorrect ones(t=2.69 p<0.03 dF=4 N=5, paired t-test) and is above chance (t=2.69 p>0.03 dF=4 N=5, one sample t-test). This we interpret as an evidence for the mice retaining the order-place association memory even at a remote time point through not being able to retrieve it immediately after a gap of 30 days.

**Figure 6:**
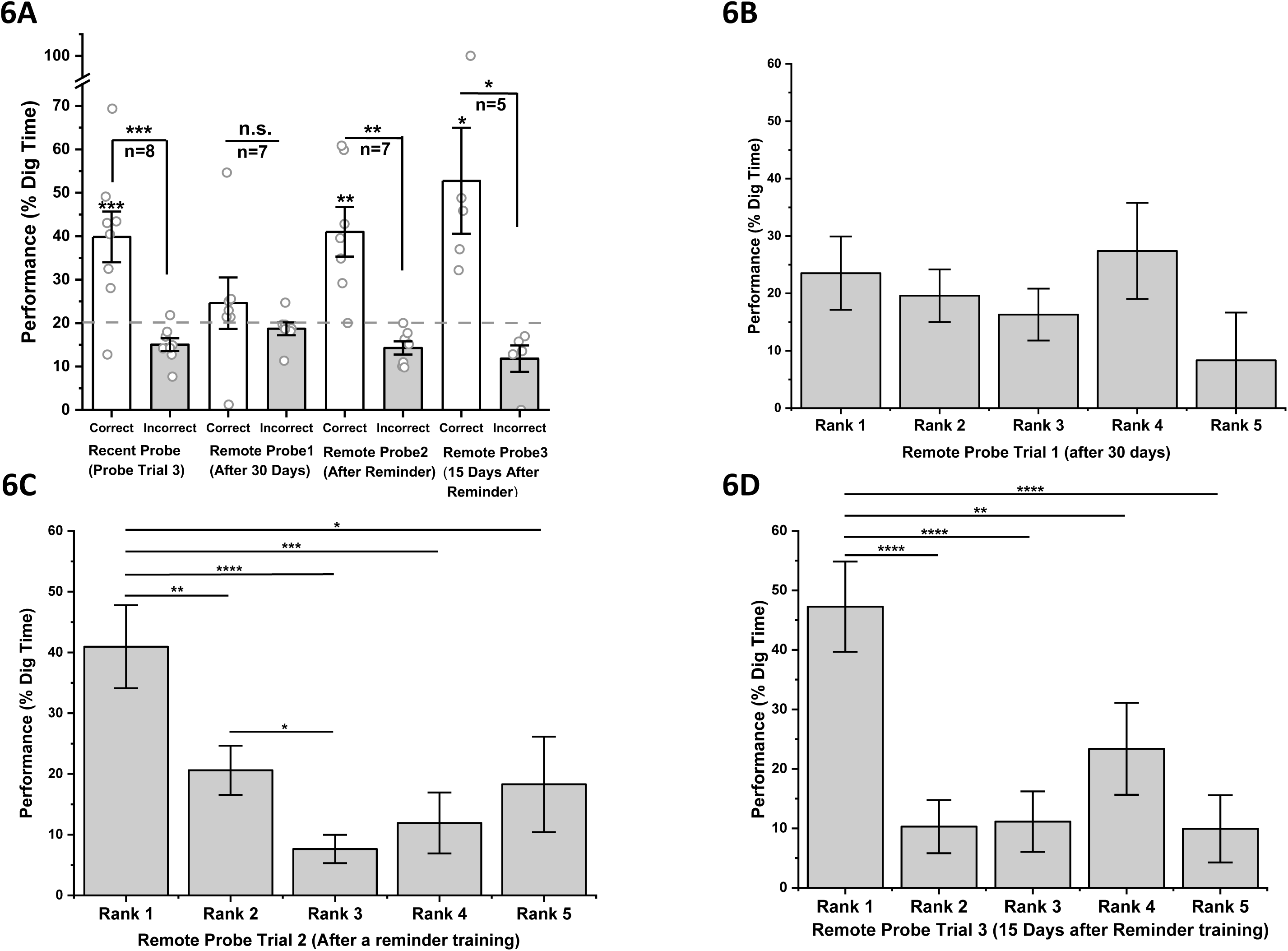
Retrieval of remote temporal order memory in the 5 Order-place association paradigm. **6A)**Open circles are the mean % dig-time for a given mouse in the correct (open/white bars) and incorrect (grey bars) locations. The mean % dig-time for the incorrect and correct locations are significantly high when probed at the recent probe, PT3, immediately after 21 training sessions (T=3.39 p<0.006 dF=7 N=8, paired t-test) but not significantly different at the remote probe trial, RPT1 conducted 30 days after the last training session(T =0.80 p=0.23 dF=6 N=7, paired t-test). However, in another probe trial RPT2, conducted after a single reminder training session, the % dig-time at the correct location is again significantly higher (t= 3.72 p<0.005 dF=6 N=7, paired sample t-test). In RPT3, a remote probe trial conducted 15 days after further reminder training sessions, the % dig-time at the correct location persists in being significantly higher than that in the incorrect locations (t=2.69 p<0.03 dF=4 N=5, 1 sample t-test). The dotted grey line signifies the chance level of 20% for a 5-position task; the mice dig at the correct location significantly higher than chance at the recent probe, PT3 (T=3.39 p<0.006 dF=7 N=8, 1 sample t-test), and at the remote probe trials RPT2 (t= 3.72 p<0.005 dF=6 N=7, 1 sample t-test) and RPT3 (t=2.69 p<0.03 dF=4 N=5, 1 sample t-test). **6B-D)** The grey bars in panels 6B,6C and 6D represent the mean % dig-time across ranks (1 to 5) measured during the remote probe trials RPT1, RPT2 and RPT3 in that order. One-way ANNOVA performed at each of the probe trials revealed that the mean % Digging across the ranks is not different for RPT1 (F=0.88 p=0.48 dF=4 N=7) but it is different for RPT2 (F=6.87 p<0.0001 dF=4 N=7) and RPT3(F=6.90 p<0.0001 dF=4 N=7). Post-hoc analysis showed that Rank 1 is higher than all other ranks in both RPT2(Fischer LSD post-hoc Rank1>2 p<0.002, Rank1>3 p<0.0001, Rank1>4 p<0.0002, Rank1>5 p<0.016, Rank2>3 p<0.033) and RPT3(Fischer LSD post-hoc Rank1>2 p<0.0001, Rank1>3 p<0.0001, Rank1>4 p<0.011, Rank1>5 p<0.001). The degree of significance of the means comparison are indicated as * with each * corresponding to an order of magnitude. * p<0.05 ** p<0.01 *** p<0.001 ****p<0.0001

Having, shown that the temporal order memory persists in these animals, we wanted to ask if the distinct signatures of higher order associations are also present. We probe this using similar analysis that is performed on the recent memory and as described above. First by dissociating the errors into ranks followed by segregating forward and backward errors following which ranking them according to proximity in temporal sequence.

Ranking of errors during remote retrieval: Fig 6b, 6c and 6d shows the percentage digging segregated into ranks for all remote probes (RPT1, RPT2 and RPT3). In accordance with our earlier description, we do not see significantly elevated digging in any rank positions in RPT1 (conducted before a reminder training). On the other hand, an ANOVA performed on the percent digging as function of ranks observed at RPT2 and RPT3 (conducted after reminder trainings) showed a main effect (F=6.87 p<0.0001 and F=6.90 p<0.0001 respectively). Post hoc analysis shows a significantly increased digging at the Rank 1 position or the correct location (RPT2 Fisher LSD post-hoc Rank1>2 p<0.002, Rank1>3 p<0.0001, Rank1>4 p<0.0002, Rank1>5 p<0.016 and in RPT3 Fisher LSD post-hoc Rank1>2 p<0.0001 Rank1>3 p<0.0001 Rank1>4 p<0.011 Rank1>5 p<0.001) indicating the animal has retrieved the memory for the correct location. However, in RPT2 specifically (the Probe trial conducted immediately after the reminder training session) the analysis also showed that Rank2 digging is significantly more than Rank3 (Fisher LSD post-hoc Rank2>3 p<0.033). The difference between Ranks 3 and above are not significant. Thus, RPT2 in the remote time mirrors the relationship between ranks in recent probe, PT3. This Rank2 elevation is not seen in RPT3.

### Forward and backward errors during remote memory recall

Differences emerge when the ranks of the recent and remote probes are dissociated into forward and backward errors. We had previously observed that as the training progresses the mice make more forward errors. This hints at the development of elimination as a strategy where they understand that the correct location cannot be one from which they have already obtained/recovered the pellet during that session. Fig. 7i shows the percent digging in incorrect locations segregated in forward and backward positions. The forward errors represented as dark grey bars are significantly higher (T=3.93 p<0002 dF=122, t-test) than the backward errors in the probe trial (PT3) conducted immediately after the training. However when tested remotely after a gap of 30 days we find a loss of this effect and it even being reversed (that is forward errors are significantly lower than the backward errors) for RPT2 (T=-2.02 dF=126 p<0.05, t-test). Now we dissociate the forward errors according to the ranks. ANOVA performed on the forward ranks of PT3 (recent probe trial) RPT2 and RTP3 (remote probes) exhibit a main effect (F=5.60 p<0.0005 and F=6.85 p<0.0001 and F=9.42 p<0.0001 respectively). In PT3 the post-hoc tests show Rank 1 to be significantly elevated above Rank 3+ and Rank4+ but not Rank2+. Rank 2+ is significantly higher than Rank3+ (as stated earlier). In the remote probes (RPT2 and RPT3) this elevation in Rank2+ is not observed in post-hoc tests, even though Rank1 is higher than all the subsequent/larger ranks RPT2, Fisher LSD post hoc Rank1>2+ p<0.0006, Rank1>3+ p<0.0004, Rank1>4+ p<0.0002, Rank1>5+ p<0.005 and in RPT3 Fisher LSD post hoc Rank1>2+ p<0.0001, Rank1>3+ p<0.0001, Rank1>4+ p<0.012 Rank1>5+ p<0.006). The rank-wise backward errors exhibit no significant differences. Elevated Rank 2+ positions in the recent probe (PT3) indicate that the mice have acquired the temporal order in the sequence beyond one-on-one associations. These ranks are indicative of inferred relationship between consecutive positions without having been trained to do so explicitly. This indicates the possibility of a transitive inference of a larger sequence order after being trained in pair-wise order-position associations. This inference is lost in the remote time point.

**Figure 7:**
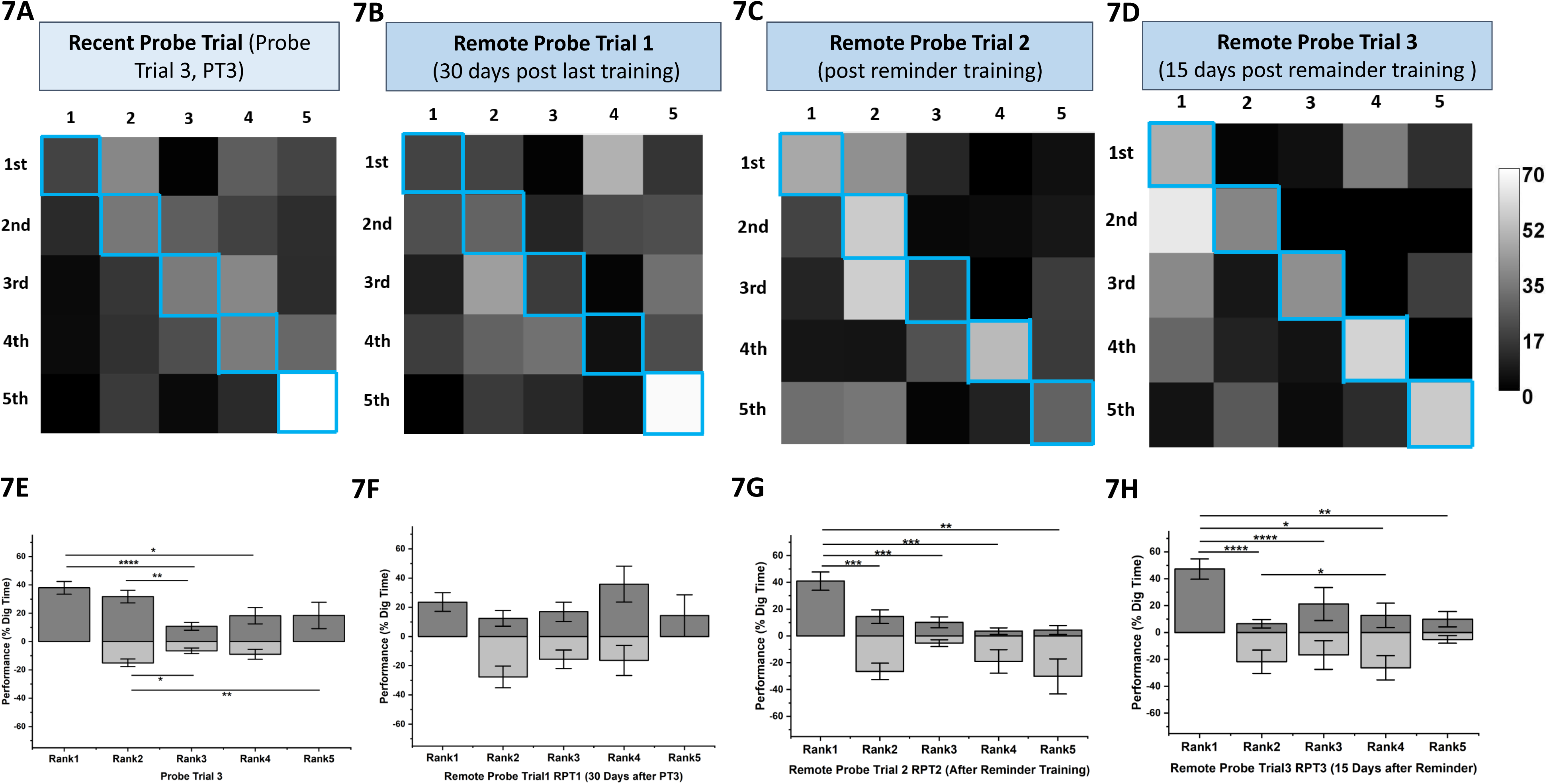
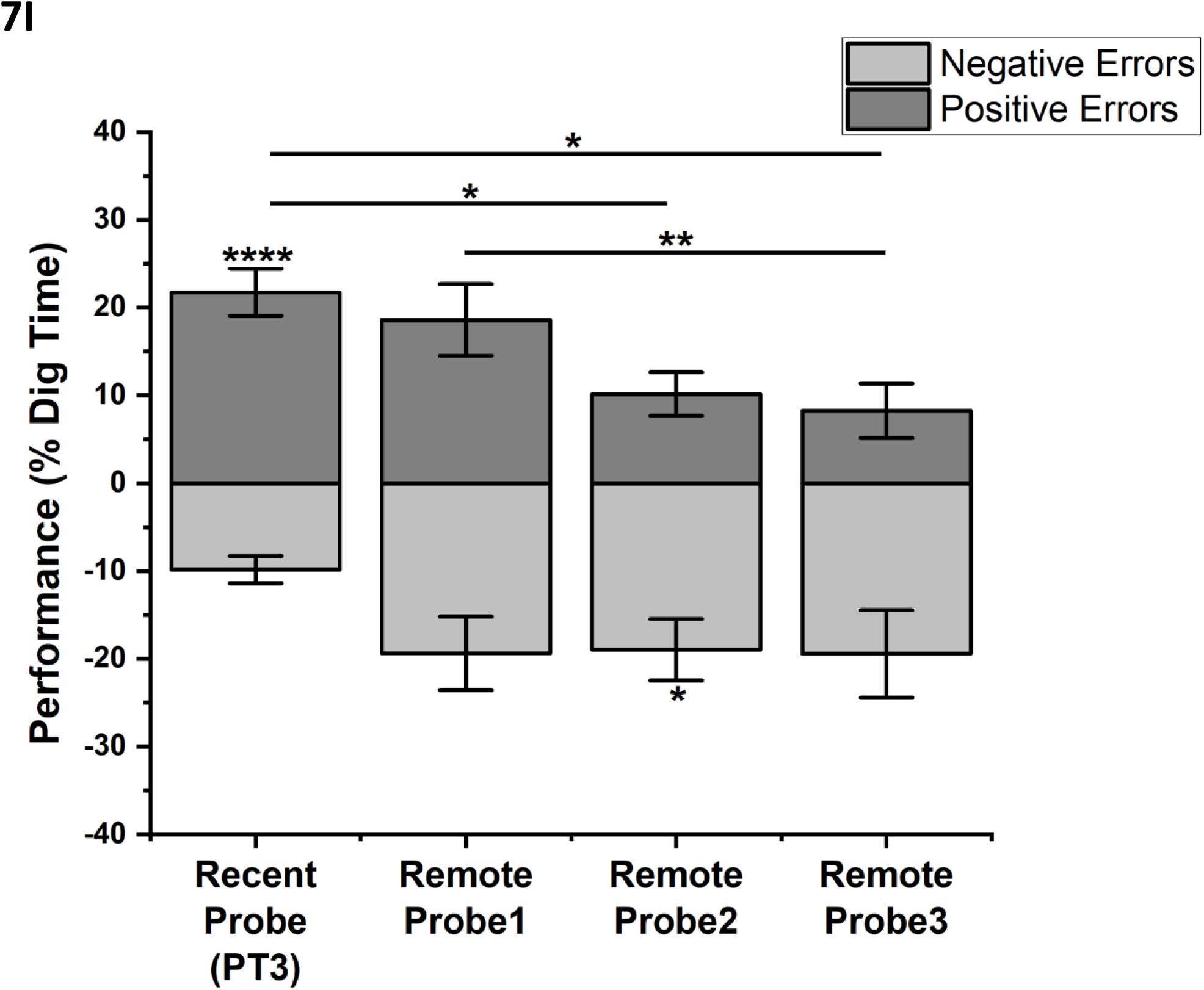
Comparison of Recent and Remote Probe trials in OPA paradigm. **7A-D)** Heatmaps comparing Recent and Remote probes. PT3, RPT2, RPT3 show concentrated digging along the diagonal/correct positions. RPT1 shows non-specific digging. The recent probe PT3 shows more diffused digging, and more forward/positive errors compared to the remote probes RPT2 and RPT3. The remote probes show less digging in the forward/positive errors and more concentrated digging along the diagonal. The diagonal elements, outlined in blue, represent the correct locations. The scale bar shows the lightest/highest % dig-time represented in the Recent(PT3) and Remote(RPT1.RPT2,RPT3) probe heatmaps to be around 70% and the darkest/lowest % dig-time to be 0%. **7E-H)** The dark grey bars represent mean % dig time in locations representing Rank1 and Rank-wise forward errors and the light grey bars represent Rank-wise backward errors across Recent (PT3, 7E) and Remote(RPT1, RPT2, RPT3 in 7F,G,H respectively) Probes. One-way ANNOVA performed at each of the probe trials revealed that the mean % digging across the forward/positive ranks is different for recent probe PT3 (F= 5.60 p<0.0005 dF=4, N=8), Remote probe RPT2 (F=6.85 p<0.0001 dF=4,N=7) and Remote probe RPT3 (F=9.42 p<0.0001 dF=4, N=5) but not for RPT1(F=1.24 p=0.30 dF=4 N=7) which does not exhibit significant order memory either. Post-hoc tests in the Recent probe, PT3 Forward/positive ranks reveal mean % dig time in Rank1(the correct location) is significantly higher in subsequent forward ranks but not in the immediately proceeding Rank2+ locations(Fischer LSD post-hoc Rank1>3+ p<0.0001, Rank1>4+ p<0.0021, Rank2+>3+ p<0.011). This elevation of 2+ is markedly absent in the remote probes, as post-tests reveal that the Rank1 mean% dig-time is significantly higher than all subsequent forward ranks including Rank 2+(RPT2 Fischer LSD post-hoc Rank1>2+ p<0.0006, Rank1>3+ p<0.0004, Rank1>4+ p<0.0002, Rank1>5+ p<0.005, RPT3 Fischer LSD post-hoc Rank1>2+ p<0.0001, Rank1>3+ p<0.0001, Rank1>4+ p<0.012 Rank1>5+ p<0.006). **7I)** The dark grey bars represent mean % dig time in locations representing total forward errors and the light grey bars represent total backward errors across Recent (PT3) and Remote Probes (RPT1, RPT2, RPT3). The recent probe PT3 shows forward errors are significantly higher than backward errors (T=3.93 p<0.0001 dF=122, N=8, t-test), However by remote probes, this effect is not seen and RPT2 exhibits higher backward errors (T=-2.02 p<0.05 dF=126, N=7, t-test). One-way ANNOVA performed to compare the forward errors reveals that the recent probe, PT3 exhibits higher forward errors than those made in the remote probes (F=3.69 p<0.02 dF=3 Fischer LSD post-hoc PT3+>RPT2+ p<0.011, PT3+>RPT3+ p<0.009). The degree of significance of the means comparison are indicated as * with each * corresponding to an order of magnitude. * p<0.05 ** p<0.01 *** p<0.001 ****p<0.0001

### Higher order associations (HOA) are dominant in order place training compared to combined training and they diminish with time

We reason that, since the mice are trained in associating correct location with either order or flavor-order cues and not explicitly on the sequence of events, their ability to infer temporal relationships among the cues would be indicative of higher order associations. Thus, in order to check if the animals have acquired these associations, we devised the difference between the % digging at Rank1 and Rank2/Rank2+ positions as a measure of HOA. Lower difference with a clear selective digging at correct location would indicate such an association being stronger. Fig 8A show this measure as a bar graph after whole probe (PT3W) and order (PT3O) probe. Comparison of these differences using t-test showed that the difference is significantly lower in order trials (Rank1 – Rank2 t= 4.84 p<0.0006 dF=11 and Rank1 – Rank2+ t=3.03 p<0.006 dF=11, paired t-test). We interpret this as indicative of the animals searching more in the Ranks2 positions in order trials compared to whole probes. However, during remote retrieval in order probes we see the Rank1 – Rank2+ difference increases and is significantly different compared to recent retrieval (PT3, RPT2, RPT3 as shown in Fig.8B). We note the remote retrieval occurs after reminder training . Thus, during the remote testing even though the direct comparison of these differences did not show a main effect across the three trials, ANOVA performed on forward errors showed that this difference is significant between RPT3 and PT3 (F= 5.36 p<0.02 Fisher LSD post-hoc PT3+/15Days+ p<0.005).

**Figure 8:**
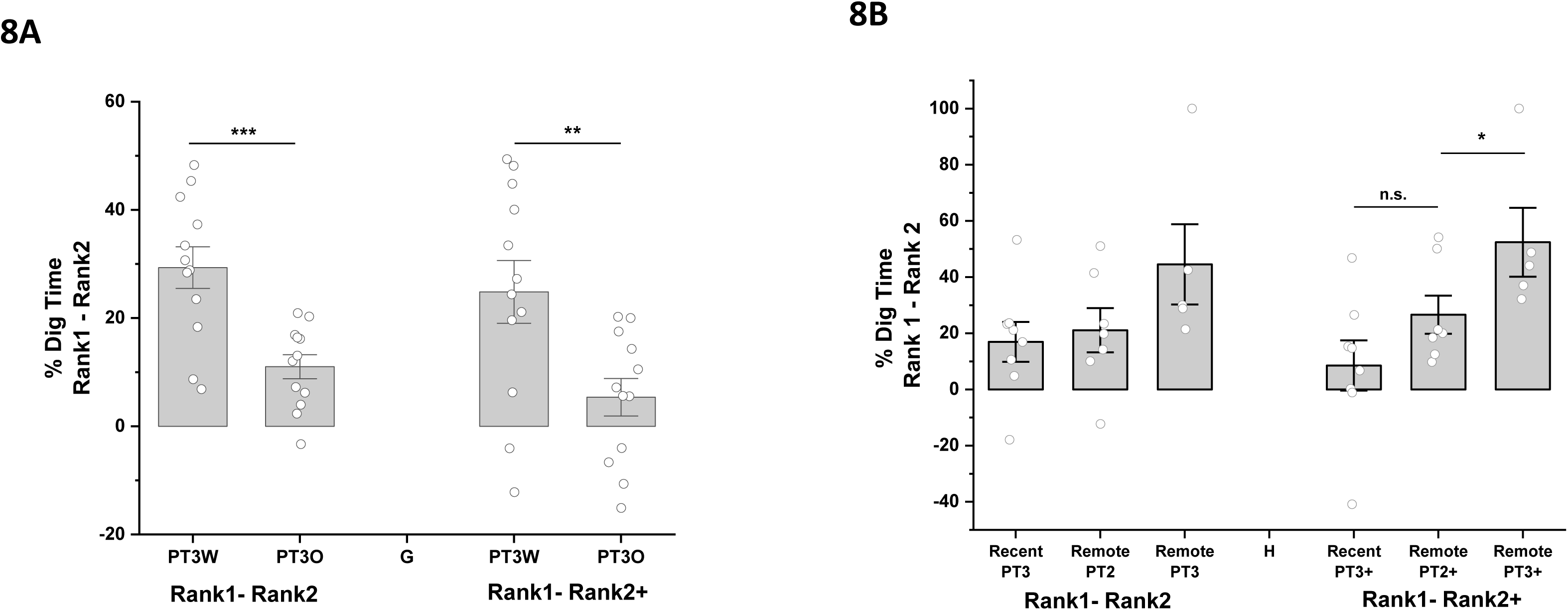
Comparison of Rank1-Rank2 differences across OPA and OFPA probes. **Fig 8A)** The open circles represents the differences between Rank1 and Rank2 / Rank1 and Rank2+ for individual mice. Grey columns represent the % mean dig-time difference between Rank1 and Rank2/Rank 2+. The difference between % dig-time in Rank1 and Rank2 positions is significantly higher in PT3W than in PT3O (T= 4.84 p<0.0006 dF=11 N=12, paired t-test). This persists even in the forward ranks, where the difference between % dig-time in Rank1 – Rank 2+ is higher in PT3W (T=3.03 p<0.006 dF=11, N=12, paired t-test). **Fig 8B)** The white circles represents the differences between Rank1 and Rank2 / Rank1 and Rank2+ for individual mice. Grey columns represent the % mean dig-time difference between Rank1 and Rank2/Rank2+ across Recent Probe trial 3 (PT3), Remote probe trial 2(RPT2 after 1 reminder session) and Remote probe trial3 (RPT3). One-way ANNOVA performed to compare mean % dig-time difference for Rank1- Rank2 across PT3, RPT2 RPT3 shows no main effect (F= 2.27 p=0.13 dF=2). However comparing the mean % dig-time for the forward/positive ranks shows an upward trend in increased Rank1-Rank2+ differences from recent to remote probes and one-way ANNOVA reveals a main effect (F=5.36 p<0.02 dF=2). The post-hoc analysis reveals significantly higher difference in the remote probe trial RPT3 compared to the recent probe trial, PT3 (Fischer LSD post-hoc p<0.005). The degree of significance of the means comparison are indicated as * with each * corresponding to an order of magnitude. * p<0.05 ** p<0.01 *** p<0.001 ****p<0.0001

## Discussion

We establish an event arena-based behavioral paradigm to test temporal order memory in rodents. In our task the mice are taught to associate their entry order with a unique location thereby making an Order Place Association memory. In this paradigm the mice receive explicit training in which they learn to dig predominately in the correct reward location pertaining to that order of entry. In any given session the mouse can dig in any of the 5 positions, each occupying a unique place in the temporal sequence of event presentation and ranking these positions according to temporal associations yields distinct signatures of mouse digging pattern.

The event-arena experiments described here, occupy a spectrum between 2-alternative-forced choice and free-recall, since the mice can utilize the previous unique position as a cue for the next location at the same time they also have a choice of all 5 positions to dig from. The temporal memory in the mice exhibits characteristic comparable to free recall studies done in humans ^42^ the mice exhibit higher forward errors than the backward errors when trained in 5 Order Place Association task as well as combined order, flavor, place association task. Given that the animals have been trained over multiple sessions spread across several days it is unlikely that this is due to recency effects. Further, our results show that the animals dig selectively in Rank 2+ positions more than in the recently exposed Rank 2- position. The mice in our task exhibit an asymmetric forward lag effect, which in human free-recall studies is attributed to recency effects and generation of temporal contexts in the hippocampus. Their capacity to make forward 0lag transitions or 2+ errors in our study could possibly be a culmination of their ability to i) Elimination Strategy : Integrate their experiences of sequence exposure across the days and during a given session to learn the rule that the correct locations they have encountered in a given day, will not be repeated ii) Higher Order Association: The increased likelihood of animal associating a given place with temporally close entry order. Therefore, the chance that a correct location could be one from the 2+ options is more likely than 2- options.

This form of learning is evident when tested in the recent time in the OPA paradigm and is lost in the remote time. This could indicate a loss of higher order association and hence the temporal context. These higher order effects/temporal inferences exist only in the recent probes and not in the remote probes suggesting that they emerge from understanding the detailed structure of the task. A single re-training session is able to serve as a reminder for only the direct pair-wise associations between A and B, B and C and so on, as is evidenced by the probe trials. However, the other details pertaining to the task structure are lost.

Similar form of learning is also observed during a combined order-flavor place association training when specifically tested for order-place association and absent when probed for flavour-place association. Had the mice exhibited higher 2+ (0 lag forward asymmetry effect) digging even in the flavor-order cued probes, this would mirror the results of list recall studies^42–44^.

The Hpc has been shown to have a role in temporal sequence memory and in transitive inference ^26^ however our efforts to lesion out the Hpc to test for its role was not fruitful. We attribute this to complexity of the task and probe structure.

## Materials and methods

### Mice, strain, housing, marking

We used male inbred C57BL/6J mice from the colony maintained in the Central Animal Facility, Indian Institute for Science. They were group housed in pairs, immediately after being weaned. The mice are housed in a holding room with a 12-hour light/dark cycle. The experiments were conducted during the light ON period, between 7 am – 6 pm. The group-housed mice were ear-marked so we could differentiate between them. All the animal behaviour was performed in accordance with the animal protocol approved by the Institute Animal Ethics Committee, Indian Institute for Science, Bengaluru.

### Arena build and Enclosure

The event arena has a square base with raised walls on the sides. In the middle of each side, is a start box with two sliding doors, one leading to the inside of the arena, and the other leading to the outside. This arena is raised onto a sturdy metal frame, for the ease of the experimenter. This set-up is enclosed in a large cuboidal chamber, made of plywood, built into which are the light panels on the top for illumination, fans for ventilation on the sides and 4 doors on 4 sides. The doors are aligned with the start boxes. The event-arena could only be accessed by opening these doors. This enclosure was built to eliminate any extra-maze cues that could be provided by the architecture of the room, the arrangements of objects in it, or even the experimenter. The camera used for observing and recording the mice is mounted directly above on the roof of the enclosure.

The square base was made of packaging pinewood of the dimensions 104 × 104 cm. The side walls were raised to a height of 15 cm and made of 2 mm acrylic sheets. The start boxes were made of Perspex sheets. The floor of the square base had a uniformly spaced 7×7 grid of drilled holes. These are used to accommodate the cups which constitute the sand-wells. This grid permitted the experimenter the flexibility to create configurations/position maps of different types. Only the holes, which marked the positions for the task had cups fitted in them, as sand-wells, while the rest are covered up. The whole base of the arena is filled with sand, usually 2-3 cm deep and it is usually smoothened out so as to eliminate lumping, height variations, patterned undulations or irregularities of any sort.

### Map of the Arena

The positions which are chosen to constitute the map of the arena, are marked using a raised ring of white acrylic surrounding the sand-well. These are visually salient enough for the mice to recognize from a distance. The two objects which are used as intra-maze cues are fixed to the floor of the arena with double sided tape to ensure they are not toppled over or displaced by the mice. One object was composed of orange ping-pong balls forming a pyramidal shape. The second object was shaped as a tower with a cylindrical bottom with a conical top, painted in a black and white pattern. An even layer of construction sand covers the entire base and only the acrylic rings around the relevant sand-wells and the visual cues are visible. The mice are handled for 6-7 days, twice a day, to reduce their anxiety.

### Habituation

The mice were habituated to the event arena to make them familiar to the experimental set-up and to a space much larger than their home cage. The objective is to reduce wall-hugging behavior and to make them explore the interior, open space of the arena, where eventually the locations with buried pellets will be present. During habituation, the two visual cues are present along with the sand-well locations, however there are no food pellets buried in any of the locations yet.

### Food restriction

The mice are provided with ad libitum food pellets (NutriLab Rodent Feed, Provimi) in their home cage, till the handling phase. Subsequently the food was restricted in a graded manner. For the first 3 days, the cage was provided with two pellets per mouse per day. For the next 2 days the animal was only provided with 1 food pellet per day. This was performed to acclimatize them to the possibility of food restriction, and to ensure their motivation for Shaping, which requires them to retrieve food pellets from the sand-wells.

Before a shaping/training/probe trial session, the animals would only be fed 1 pellet. Whereas on ‘rest’ days, when they do not have to perform in the arena the next day, they would be fed 2 pellets each. This dietary regimen was stressful for the mice, and it was ensured that the mice do not undergo more than 2 consecutive days of training /food restriction. The mice were therefore continuously monitored, if a mouse lost weight /appeared weak they are provided with food and rested from performing the task. If the two cage mates used to fight and snatch each other pellets, they were individually fed and monitored, so that a dominant animal did not starve out it’s meeker cage-mate.

### Shaping

Digging and burrowing are natural behaviors for mice. However, to perform the task the mice have to know of and be able to dig out a pellet buried 2-3 centimetres deep in a sand filled cup. They need to be made aware of the ‘rule’ that only the sand-wells in the 4/5 marked positions have at some depth contain the food-reward. To make them pay attention to the fact that these specific locations in the arena are the ones which will have the pellet buried in them, we placed one food pellet on each of the locations such that they are above the sand surface and visible. Over successive shapping sessions we buried them little by litte into the sand such that at the end 6 sessions the pellet is buried completely inside the sand well. At the end of the shaping session the mice were able to dig to the appropriate depth to retrieve these pellets.

The entry of the mouse into the arena was randomized, and in a day, they would have entered through each of the 4 start boxes in no particular order. This is necessary for the mice to use allocentric navigation to form a map of the arena. Every sand-well base had the same mixture of 5 flavours crushed and placed under a mesh. However, for the purpose of shaping, unflavoured food pellets are used as food rewards, since all locations contained the food reward. This is done to ensure that no one location to be associated with any particular flavor.

### Flavor pellets

The non-flavoured and flavoured food pellets used outside the arena as cues, and inside as buried food rewards are custom prepared.

The rodent food pellets (NutriLab Rodent Feed, Provimi) are ground into a semi-course powder in a kitchen grinder. The powdered food is filtered through a 1.5 mm sieve size mesh, to remove any large chunks. The powder is then evenly mixed with the flavor (flavor concentrations in w/w are mentioned below) in the appropriate amounts. Out of the 10 flavors used, 9 were in solid powdered form, so they are mixed with the powdered food directly. 1 of those flavors was available only in liquid form, and is thus, mixed with the water which is used to form the dough.

### Food flavours

a. Basil (Snapin, Duval enterprises Pvt. Ltd.) – 1% w/w in powdered mouse feed.
b. Cumin (Everest, S. Narendrakumar & Co.) – 1% w/w in powdered mouse feed.
c. Dry Mango (Everest, S. Narendrakumar & Co.) – 1% w/w in powdered mouse feed.
d. Ginger (Snapin, Duval enterprises Pvt. Ltd.) – 1% w/w in powdered mouse feed.
e. Cocoa (WeikField, Weikfield foods Pvt. Ltd.) – 1% w/w in powdered mouse feed.
f. Thyme (Keya, Keya Foods International Pvt. Ltd.) – 0.5% w/w in powdered mouse feed.
g. Cassia (Arvind Trading Company) – 0.5% w/w in powdered mouse feed.
h. Elaichi/Cardamom (Arvind Trading Company) – 1% w/w in powdered mouse feed.
i. Rosewhite liquid essence (Bush, International Flavours and fragrances India Pvt. Ltd.) – 1% volume/w in powdered food.
j. Fenugreek (Eastern, Eastern Condiments Pvt. Ltd.) – 1% w/w in powdered mouse feed.

Note: Source of all the condiments is kept the same throughout the experiments.

### Composition and dimensions of sandwells

The sand wells were composed of the plastic cups of a standard size available readily in the market. The top of the cup has a protruding rim which rests on the surface of base of the event arena, while the rest of the cup juts out below the surface of the arena. The bottom of the cup is fitted with a sponge and above it the crushed flavoured pellets. A fine mesh (0.5 mm sieve size) fits atop the crushed pellets. The reward pellet is kept on top of the mesh, and sand is filled throughout the cavity of the cup, from top to the mesh. The mesh is too fine to allow sand to enter below, into the cavity containing the crushed flavoured pellets. This is necessary, as the crushed flavoured pellets form the background odour of every sand-well.

### Training protocol

Training is performed to make the mice associate one unique location to a particular position within a temporal sequence. 1 training session is spread over about 3.5 hours. Mice undergo only 1 training session in a day, as the task is spread over a long period of time and is strenuous. Each training session has 4/5 entry events. A sheet with the randomized flavor presentation order, and entry-box order is prepared before hand.

The mice cages are brought from the holding room to the room containing the behaviour set-up 45 minutes before the task commences. We utilize this time to ready the cups which will be used as sand-wells. The bottom of each cup (under the mesh) contains roughly crushed pellets, 2 of each flavor. The cups that constitute the sand-wells containing the reward contain all the crushed pellets except those flavoured ones which will be kept above the mesh, accessible for the mice to dig and retrieve. These cups and crushed pellets are prepared fresh for each training sessions.

When the task begins with the 1^st^ entry event, when the mice would be allowed to feed on a flavoured pellet in their homecage for 30-40 seconds. The pellet is then taken from them, and they are let into the arena via the entry-box. It is important to ensure they aren’t holding on to the cued food pellet, since we have observed that even a small pellet when they hold on to, the mice gets pre-occupied with them in the arena, and they will not dig for food. We start recording the video upon entry and we monitor their behaviour for 2 minutes, during which they are free to explore and dig the sand-wells. The trial ends, and we stop recording once the 2 minutes are over, or when the mouse has dug out and found the food pellet and is consuming it. If the mouse has not found the pellet during this time period, the experimenter directs the mouse to the correct well, with their finger placed on and gently scratching the top of the correct well. Over time, as the training sessions progress, the mice ‘learn’ that the experimenter keeping a finger on a sand-well indicates that it’s a reward containing well and that they can find the food pellet upon digging there. After the mouse has dug and retrieved the pellet, it has 10-15 seconds to nibble on it, after which we remove the mouse from the event arena, take away the remaining pellet and place it back into its home-cage. It is important to take away the pellet from the animal, before placing it into the home cage with its cage-mate as this could cause a fight to break out between the two mice, since both animals are food-deprived.

The sand is shuffled around and smoothed after every trial to erase footprints or obfuscate any left-over odor cues strewn around the wells. The cups are removed, and fresh cups are put in place. The 2 prepared copies of the cups are shuffled alternately every trial. 70% Alcohol is sprayed onto the sand and is concentrated a little more inside the sand-wells themselves, to ensure a thorough obfuscation of odor-cues coming from the pellets in the sand wells. This step is extremely important to ensure that this task is primarily a visual cue-guided spatial navigational task, and not an odour directed one. If the mice can see (if the pellets are sticking out accidentally) or can sniff out (if the reward well was the only well containing any flavor smell) the correct well, they have no incentive to learn the navigational map in accordance with the order (or falvor) cues. It ceases to be a multiple-association memory task. We have already ensured the reward well does not contain any extra odor, over and above the other non-reward wells, by means of the crushed pellets on the base of the cups, under the mesh.

After preparing the event-arena and the cups for the next entry, we cue the next mouse with flavoured pellets in its home cage. The Entry 1 (corresponding to order 1, position 1) is performed for all the mice in the batch, before moving on to Entry 2 (corresponding to order 2, position 2) for the same mice, in the same order. This ensures that each mouse has a gap of 45 minutes before each entry. 45 minutes after all the sessions are done for the training day, the mouse-cages are kept back in the holding room.

### Probe trial protocol

The strength of the memory is measured using Probe trials. It was critical that the animals do not sense any change in the experimental set-up, experimenter, or protocol when the probe trial is being conducted. These are conducted similar to the training session with crucial differences; 1. We do not place a buried food pellet at the bottom of the sand well 2. We leave the mice in the arena for 60 seconds (and not 120 seconds) to monitor its digging pattern. To ensure the probe trial does not lead to extinction, the experimenter at the end of the session, discretely buries the pellet at the correct location and leads the mouse to it. The stronger the association made between the order and location the longer they will dig at the correct location. The percentage digging time at the correct location is considered as a measure of their memory, with 25% (for 4 position event arena) and 20% (for 5 position event arena) dig time at one particular location is considered chance performance. Performance is measured as the percentage ratio of digging time at one particular location to the total digging time in all locations for that trial.

For OFPA 5-position paradigm, to reduce the extinction effects we had split all the probe trails across two days with two of the flavours/entry orders being probed on one day and the remaining on the other day. Whole probes were conducted first, and order probes immediately after. The schematic detailing the structure of the order probe is provided in Figure 4B.

### HeatMap generation for visualization

Fig 3A shows a schematic illustration of a 5 × 5 matrix, where the rows represent the order of entry into the arena and the columns represent the locations/positions in the temporal sequence. The 5 diagonal elements represent the correct location, and every other location on either side of the diagonal (the remaining 20 elements/locations) corresponds to an error. The set of errors outlined in yellow correspond to forward/positive errors, where the mouse digs in positions proceeding those in the temporal order (and those positions which it has not yet encountered in the current training/probing session). The set of errors outlined in orange correspond to backward/negative errors, where the mouse digs in positions preceding those in the temporal order (and those which it has encountered before in the current training/probing session). This example figure shows the ideal heatmap, where the mouse has learnt the order-place associations. The diagonal shows the highest amount of digging. The positions adjacent to the correct location show reduced digging in a graded way, depending on the temporal distance from the correct location. In this example figure, the digging reduces in the incorrect locations, by the same measure, in the forward/positive and backward/negative directions.

Intensity at each of the 25 elements represents the average performance (as percentage dig-time) for all mice at that position for a particular nth entry. No normalization has been performed.

Data and Code Sharing: We will be able to the share the data acquired in this study and the analysis pipelines that we have used after acceptance of the manuscript upon a reasonable written request.

## Author contributions

SS and BJ designed performed and analysed the behavioral experiments.

VPS helped in designing the event arena tasks, took part in discussions and also performed some of the experiments involving four position OPA task.

RB and SK both helped in performing the four position OPA task.

SS and BJ wrote the manuscript.

## Funding

This work is funded by grants to BJ from SERB (EMR/2017/004155), DBT Grant DBTO/BCN/BJ/0402, DSTO DSTO/BCN/BJ/1102, DBT IISc Partnership, Ramanujan Fellowship, Tata Trust JTT/MUM/INST/IIOS/201314/0033 and Pratiksha Trust.

SS received funding from CSIR (CSIR 09/079/(2624)/2013-EMR-I) .

SK was supported by Tata trusts JTT/MUM/INST/IIOS/201314/0033.

## Supplementary Figures

**Supplementary Figure 1:**
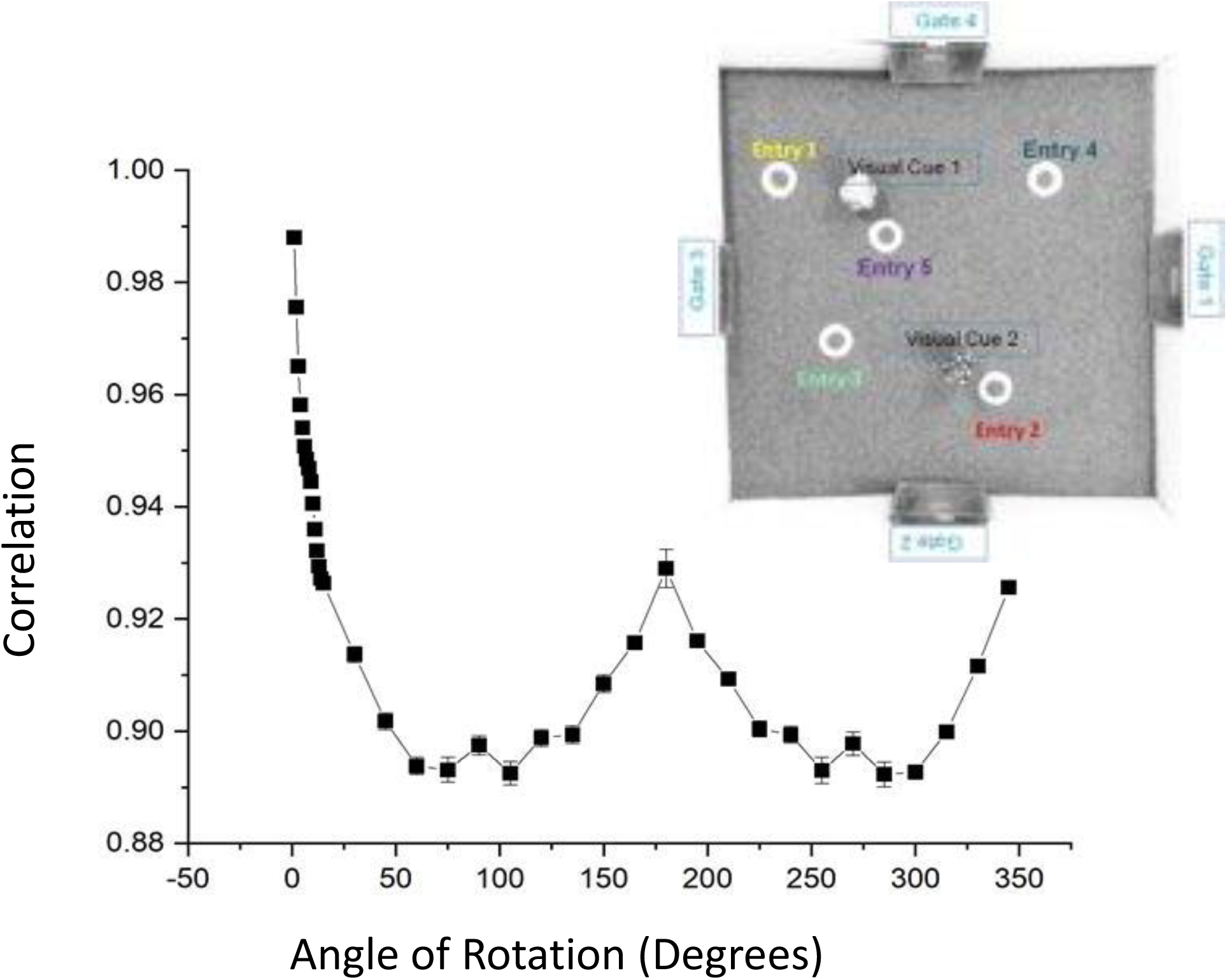
Image correlation provides a quantitative estimate of the rotational symmetry in 5 position Event-Arena. The top view of the event arena is rotated in increments of 1 degree initially (till 15 degrees) and subsequently by 15 degrees and the image correlation across this sequence of images is measured using Imagej (using plugin Image CorrelationJ 1o). The correlation (black squares) measures the similarity and hence the symmetry in our set-up.

**Supplementary Figure 2:**
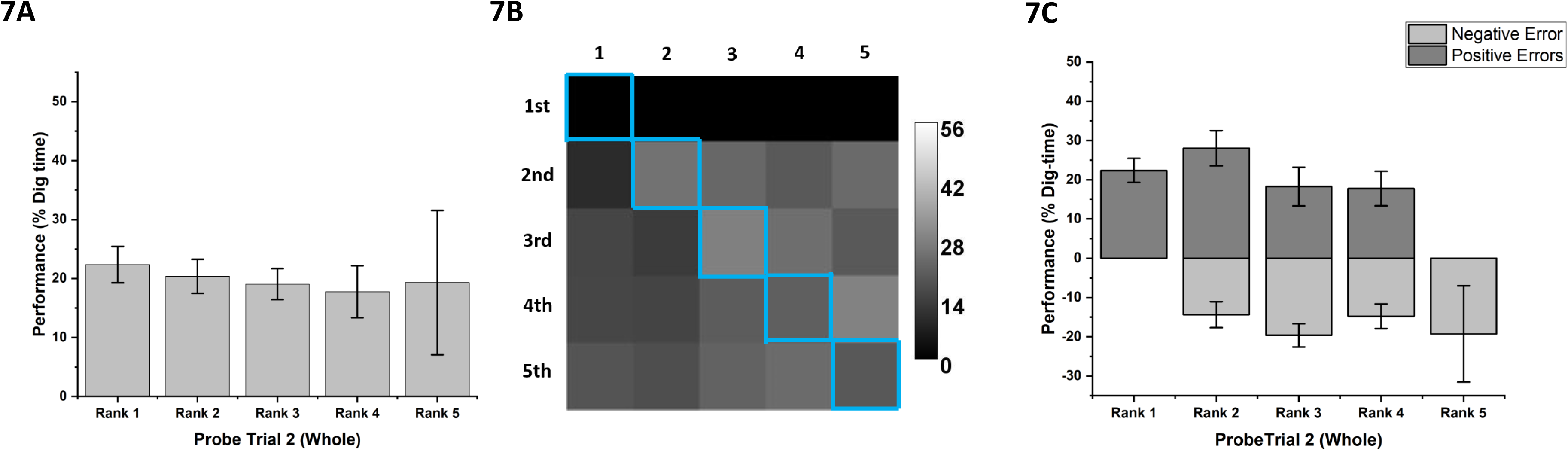
PT2W of the Combined Flavor-order place association event arena task. In the OFPA paradigm three probe trials were conducted after 3 (PT1), a total of 15 (PT2) and a total of 21 (PT3) sessions. Figure 4A shows the % dig-time in correct and incorrect locations in PT2W (PT2O, for order was not conducted), which was not significant(T=0.94 p=0.19 dF=5, N=6). PT2W was distributed across two days, and the first flavor-order pair was not probed for on either days. The grey bars in panel 2A represent the mean % dig-time across ranks (1 to 5) measured during the second probe trial, PT2W. One-way ANNOVA performed at each of the probe trials revealed that the mean % Digging across the ranks is not different (F=0.21 p=0.93 dF=4, N=6). The heatmap in 2B shows a diffused digging pattern, which is not concentrated around the diagonal/correct locations. The scale bar shows the lightest/highest % dig-time to be around 56% and the darkest/lowest % dig-time to be 0%. The first row, corresponding to Position1 and the flavor Elaichi/Cardamom, is blank because the mice are not probed for it. The dark grey bars represent mean % dig time in locations representing Rank1 and Rank-wise forward errors and the light grey bars represent Rank-wise backward errors of PT2W in 2C. One-way ANNOVA revealed that the mean % digging across the forward/positive ranks is not different (F= 0.73 p=0.54 dF=4, N=6). The degree of significance of the means comparison are indicated as * with each * corresponding to an order of magnitude. * p<0.05 ** p<0.01 *** p<0.001 ****p<0.0001

**Supplementary Table 1:**

1.1 Comparison of Correct vs Incorrect % dig-time
|  |  |  |  |  |  | T- Test (C vs Inc) |  |  | N |
| --- | --- | --- | --- | --- | --- | --- | --- | --- | --- |
|  |  |  | Mean | SD | SEM | T | dF | Prob>t |  |
| OPA (4 position) | PT1 | Correct | 28.63319 | 11.46942 | 3.82314 | 0.95032 | 8 | 0.18489 | 9 |
|  |  | Incorrect | 23.78894 | 3.82314 | 1.27438 |  |  |  |  |
|  | PT2 | Correct | 32.45769 | 8.22418 | 2.90769 | 2.56482 | 7 | 0.01864 | 8 |
|  |  | Incorrect | 22.5141 | 2.74139 | 0.96923 |  |  |  |  |
| OPA (5 Position) | PT1 | Correct | 23.13628 | 19.30397 | 7.88081 | 0.39796 | 5 | 0.35354 | 6 |
|  |  | Incorrect | 19.21593 | 4.82599 | 1.9702 |  |  |  |  |
|  | PT2 | Correct | 35.45105 | 32.72359 | 11.56954 | 1.33549 | 7 | 0.11175 | 8 |
|  |  | Incorrect | 16.13724 | 8.1809 | 2.89238 |  |  |  |  |
|  | PT2 | Correct | 39.82946 | 16.50377 | 5.83496 | 3.39839 | 7 | 0.00573 | 8 |
|  |  | Incorrect | 15.04263 | 4.12594 | 1.45874 |  |  |  |  |
|  | RPT1 | Correct | 24.58853 | 15.66988 | 5.92266 | 0.79725 | 6 | 0.22785 | 7 |
|  |  | Incorrect | 18.69093 | 3.92074 | 1.4819 |  |  |  |  |
|  | RPT2 | Correct | 41.00493 | 15.12565 | 5.71696 | 3.71947 | 6 | 0.00493 | 7 |
|  |  | Incorrect | 14.27258 | 4.04854 | 1.53021 |  |  |  |  |
|  | RPT3 | Correct | 52.7512 | 27.24201 | 12.183 | 2.68827 | 4 | 0.02738 | 5 |
|  |  | Incorrect | 11.8122 | 6.8105 | 3.04575 |  |  |  |  |
| OFPA (5 Position) | PT1W | Correct | 16.25818 | 9.60789 | 2.66475 | -1.40419 | 13 | 0.90719 | 14 |
|  |  | Incorrect | 20.93545 | 2.40197 | 0.66619 |  |  |  |  |
|  | PT2W | Correct | 22.00206 | 5.20574 | 2.12523 | 0.94204 | 5 | 0.19471 | 6 |
|  |  | Incorrect | 19.49949 | 1.30143 | 0.53131 |  |  |  |  |
|  | PT3W | Correct | 43.69362 | 8.3391 | 2.40729 | 9.84245 | 11 | 4.33E-07 | 12 |
|  |  | Incorrect | 14.07659 | 2.08477 | 0.60182 |  |  |  |  |
|  | PT1O | Correct | 20.67533 | 8.63266 | 2.30717 | 0.29271 | 13 | 0.38718 | 14 |
|  |  | Incorrect | 19.83117 | 2.15816 | 0.57679 |  |  |  |  |
|  | PT2O | Correct | 32.07597 | 9.48014 | 2.73668 | 4.41263 | 11 | 5.21E-04 | 12 |
|  |  | Incorrect | 16.98101 | 2.37004 | 0.68417 |  |  |  |  |

1.2 Comparison of mean % dig-time in Ranks in OPA (5-Position) Paradigm
| OPA<br>(5-position)<br>Rank Means |  |  |  |  |  |  |  |  |  |
| --- | --- | --- | --- | --- | --- | --- | --- | --- | --- |
|  |  | Mean | S.E.M |  | Mean | S.E.M |  | Mean | S.E.M |
| PT1 | Rank 1 | 24.25553 | 7.32604 | Rank1 | 24.25553 | 7.32604 |  |  |  |
|  | Rank 2 | 11.99414 | 3.97589 | Rank2+ | 6.53773 | 2.92078 | Rank2- | 17.7916 | 7.45224 |
|  | Rank 3 | 14.40318 | 5.79953 | Rank3+ | 16.16622 | 8.88403 | Rank3- | 12.6401 | 7.791 |
|  | Rank 4 | 25.17151 | 8.00924 | Rank4+ | 13.08693 | 9.2689 | Rank4- | 37.2561 | 12.136 |
|  | Rank 5 | 46.40008 | 14.08361 | Rank5+ | 53.90476 | 19.96448 | Rank5- | 37.0192 | 21.74828 |
| PT2 | Rank 1 | 33.16302 | 8.00187 | Rank1 | 33.16302 | 8.00187 |  |  |  |
|  | Rank 2 | 15.55492 | 3.97421 | Rank2+ | 14.42072 | 5.46595 | Rank2- | 16.5905 | 5.84433 |
|  | Rank 3 | 19.72997 | 5.16827 | Rank3+ | 18.3656 | 8.36124 | Rank3- | 20.7912 | 6.6978 |
|  | Rank 4 | 18.24018 | 6.09119 | Rank4+ | 21.56309 | 9.60312 | Rank4- | 15.9397 | 8.13877 |
|  | Rank 5 | 8.75389 | 6.31154 | Rank5+ | 1.04079 | 0.61837 | Rank5- | 13.896 | 10.30209 |
| PT3 | Rank 1 | 37.98623 | 4.49821 | Rank1 | 37.98623 | 4.49821 |  |  |  |
|  | Rank 2 | 23.27533 | 2.8436 | Rank2+ | 31.80476 | 4.50779 | Rank2- | 15.074 | 2.73432 |
|  | Rank 3 | 8.55612 | 1.68586 | Rank3+ | 10.81203 | 2.77412 | Rank3- | 6.52579 | 1.96488 |
|  | Rank 4 | 13.24615 | 3.35334 | Rank4+ | 18.28855 | 5.80914 | Rank4- | 8.9795 | 3.56153 |
|  | Rank 5 | 8.39498 | 4.92525 | Rank5+ | 18.46895 | 9.32232 | Rank5- | 0 | 0 |
| RPT1 | Rank 1 | 23.52736 | 6.4023 | Rank1 | 23.52736 | 6.4023 |  |  |  |
|  | Rank 2 | 19.6047 | 4.56198 | Rank2+ | 12.42251 | 5.2874 | Rank2- | 27.7237 | 7.42818 |
|  | Rank 3 | 16.30822 | 4.52058 | Rank3+ | 16.92129 | 6.54436 | Rank3- | 15.587 | 6.3388 |
|  | Rank 4 | 27.3976 | 8.37181 | Rank4+ | 35.87652 | 12.32099 | Rank4- | 16.375 | 10.34551 |
|  | Rank 5 | 8.33333 | 8.33333 | Rank5+ | 14.28571 | 14.28571 | Rank5- | 0 | 0 |
| RPT2 | Rank 1 | 40.94761 | 6.83266 | Rank1 | 40.94761 | 6.83266 |  |  |  |
|  | Rank 2 | 20.6104 | 4.04409 | Rank2+ | 14.54079 | 5.0647 | Rank2- | 26.4466 | 6.14029 |
|  | Rank 3 | 7.65111 | 2.33539 | Rank3+ | 10.2427 | 4.02545 | Rank3- | 5.31868 | 2.55476 |
|  | Rank 4 | 11.92686 | 5.01194 | Rank4+ | 3.66667 | 2.48429 | Rank4- | 19.007 | 8.76822 |
|  | Rank 5 | 18.28505 | 7.86675 | Rank5+ | 4.44444 | 3.29609 | Rank5- | 30.1484 | 13.08735 |
| RPT3 | Rank 1 | 47.25771 | 7.58968 | Rank1 | 47.25771 | 7.58968 |  |  |  |
|  | Rank 2 | 10.29012 | 4.45696 | Rank2+ | 0.5102 | 0.5102 | Rank2- | 21.7 | 8.68789 |
|  | Rank 3 | 11.1353 | 5.07419 | Rank3+ | 6.59216 | 3.11954 | Rank3- | 16.688 | 10.67773 |
|  | Rank 4 | 23.36397 | 7.73775 | Rank4+ | 21.2294 | 12.23785 | Rank4- | 26.2101 | 8.99068 |
|  | Rank 5 | 9.90927 | 5.6556 | Rank5+ | 12.8022 | 9.02655 | Rank5- | 5.08772 | 2.88808 |

| OPA (5-position)<br>Rank<br>Analysis | 1 way-ANOVA RESULTS |  |  |  |  |
| --- | --- | --- | --- | --- | --- |
|  |  | F Value | Prob>F | DF | N |
| PT1 | Total Ranks | 2.75697 | 0.03192 | 4 | 6 |
|  | Positive Ranks | 2.86222 | 0.03096 | 4 | 6 |
|  | Negative Ranks | 1.026 | 0.40176 | 3 | 6 |
| PT2 | Total Ranks | 1.83268 | 0.12638 | 4 | 8 |
|  | Positive Ranks | 1.48317 | 0.2166 | 4 | 8 |
|  | Negative Ranks | 0.13556 | 0.93842 | 3 | 8 |
| PT3 | Total Ranks | 12.99633 | 4.12E-09 | 4 | 8 |
|  | Positive Ranks | 5.59724 | 4.77E-04 | 4 | 8 |
|  | Negative Ranks | 3.72732 | 0.01582 | 3 | 8 |
| RPT1 | Total Ranks | 0.88055 | 0.47726 | 4 | 7 |
|  | Positive Ranks | 1.23555 | 0.30143 | 4 | 7 |
|  | Negative Ranks | 1.33737 | 0.27253 | 3 | 7 |
| RPT2 | Total Ranks | 6.87574 | 4.03E-05 | 4 | 7 |
|  | Positive Ranks | 6.8473 | 7.60E-05 | 4 | 7 |
|  | Negative Ranks | 2.63964 | 0.05711 | 3 | 7 |
| RPT3 | Total Ranks | 6.90538 | 7.93E-05 | 4 | 5 |
|  | Positive Ranks | 9.41712 | 9.39E-06 | 4 | 5 |
|  | Negative Ranks | 0.42916 | 0.73382 | 3 | 5 |

1.3 Means comparisons of mean % dig-time in Ranks in OPA (5-Position) Paradigm PT1 RANKS
| Fisher LSD Test | MeanDiff | SEM | t Value | Prob | Alpha | Sig | LCL | UCL |
| --- | --- | --- | --- | --- | --- | --- | --- | --- |
| Rank 2 Rank 1 | -12.2614 | 8.41028 | -1.45791 | 0.148 | 0.05 | 0 | -28.9471 | 4.42436 |
| Rank 3 Rank 1 | -9.85235 | 8.8396 | -1.11457 | 0.26771 | 0.05 | 0 | -27.3899 | 7.68517 |
| Rank 3 Rank 2 | 2.40904 | 7.90064 | 0.30492 | 0.76106 | 0.05 | 0 | -13.2656 | 18.08369 |
| Rank 4 Rank 1 | 0.91598 | 9.99796 | 0.09162 | 0.92719 | 0.05 | 0 | -18.9197 | 20.75164 |
| Rank 4 Rank 2 | 13.17737 | 9.17828 | 1.43571 | 0.1542 | 0.05 | 0 | -5.03207 | 31.38681 |
| Rank 4 Rank 3 | 10.76833 | 9.57322 | 1.12484 | 0.26335 | 0.05 | 0 | -8.22467 | 29.76132 |
| Rank 5 Rank 1 | 22.14456 | 12.00355 | 1.84483 | 0.06802 | 0.05 | 0 | -1.67015 | 45.95926 |
| Rank 5 Rank 2 | 34.40595 | 11.32991 | 3.03674 | 0.00305 | 0.05 | 1 | 11.92773 | 56.88417 |
| Rank 5 Rank 3 | 31.9969 | 11.65215 | 2.74601 | 0.00715 | 0.05 | 1 | 8.87936 | 55.11444 |
| Rank 5 Rank 4 | 21.22858 | 12.55362 | 1.69103 | 0.09395 | 0.05 | 0 | -3.67744 | 46.13459 |

PT3 Ranks
| Fisher LSD Test | MeanDiff | SEM | t Value | Prob | Alpha | Sig | LCL | UCL |
| --- | --- | --- | --- | --- | --- | --- | --- | --- |
| Rank 2 Rank 1 | -14.7109 | 4.25287 | -3.45905 | 7.06E-04 | 0.05 | 1 | -23.1142 | -6.30763 |
| Rank 3 Rank 1 | -29.4301 | 4.51953 | -6.51177 | 1.05E-09 | 0.05 | 1 | -38.3603 | -20.5 |
| Rank 3 Rank 2 | -14.7192 | 4.00183 | -3.67812 | 3.27E-04 | 0.05 | 1 | -22.6265 | -6.81197 |
| Rank 4 Rank 1 | -24.7401 | 5.07733 | -4.87265 | 2.77E-06 | 0.05 | 1 | -34.7724 | -14.7078 |
| Rank 4 Rank 2 | -10.0292 | 4.62254 | -2.16963 | 0.03161 | 0.05 | 1 | -19.1629 | -0.89548 |
| Rank 4 Rank 3 | 4.69003 | 4.869 | 0.96324 | 0.33697 | 0.05 | 0 | -4.93064 | 14.31071 |
| Rank 5 Rank 1 | -29.5913 | 6.55373 | -4.51518 | 1.27E-05 | 0.05 | 1 | -42.5408 | -16.6417 |
| Rank 5 Rank 2 | -14.8804 | 6.20805 | -2.39694 | 0.01776 | 0.05 | 1 | -27.1469 | -2.61383 |
| Rank 5 Rank 3 | -0.16114 | 6.39368 | -0.0252 | 0.97993 | 0.05 | 0 | -12.7944 | 12.47217 |
| Rank 5 Rank 4 | -4.85117 | 6.79944 | -0.71347 | 0.47667 | 0.05 | 0 | -18.2862 | 8.58387 |

**RPT2 RANKS**
| **Fisher LSD Test** | **MeanDiff** | **SEM** | **t Value** | **Prob** | **Alpha** | **Sig** | **LCL** | **UCL** |
| --- | --- | --- | --- | --- | --- | --- | --- | --- |
| **Rank 2 Rank 1** | **-20.3372** | **6.31827** | **-3.21879** | **0.00157** | **0.05** | **1** | **-32.8183** | **-7.85618** |
| **Rank 3 Rank 1** | **-33.2965** | **6.72205** | **-4.95333** | **1.89E-06** | **0.05** | **1** | **-46.5752** | **-20.0179** |
| **Rank 3 Rank 2** | **-12.9593** | **6.00396** | **-2.15846** | **0.03243** | **0.05** | **1** | **-24.8194** | **-1.09915** |
| **Rank 4 Rank 1** | **-29.0208** | **7.39727** | **-3.92317** | **1.31E-04** | **0.05** | **1** | **-43.6332** | **-14.4083** |
| Rank 4 Rank 2 | -8.68353 | 6.75139 | -1.28618 | 0.2003 | 0.05 | 0 | -22.0201 | 4.65307 |
| Rank 4 Rank 3 | 4.27576 | 7.13068 | 0.59963 | 0.54963 | 0.05 | 0 | -9.81009 | 18.36161 |
| **Rank 5 Rank 1** | **-22.6626** | **9.21465** | **-2.45941** | **0.01502** | **0.05** | **1** | **-40.8651** | **-4.46006** |
| Rank 5 Rank 2 | -2.32535 | 8.70468 | -0.26714 | 0.78972 | 0.05 | 0 | -19.5205 | 14.86976 |
| Rank 5 Rank 3 | 10.63394 | 9.00204 | 1.18128 | 0.2393 | 0.05 | 0 | -7.14857 | 28.41646 |
| Rank 5 Rank 4 | 6.35818 | 9.51685 | 0.6681 | 0.50507 | 0.05 | 0 | -12.4413 | 25.15765 |

**RPT3 RANKS**
| **Fisher LSD Test** | **MeanDiff** | **SEM** | **t Value** | **Prob** | **Alpha** | **Sig** | **LCL** | **UCL** |
| --- | --- | --- | --- | --- | --- | --- | --- | --- |
| **Rank 2 Rank 1** | **-36.9676** | **7.87925** | **-4.69177** | **1.10E-05** | **0.05** | **1** | **-52.6478** | **-21.2874** |
| **Rank 3 Rank 1** | **-36.1224** | **8.33342** | **-4.33464** | **4.21E-05** | **0.05** | **1** | **-52.7064** | **-19.5384** |
| Rank 3 Rank 2 | 0.84518 | 7.51344 | 0.11249 | 0.91072 | 0.05 | 0 | -14.1071 | 15.79741 |
| **Rank 4 Rank 1** | **-23.8937** | **9.11705** | **-2.62078** | **0.0105** | **0.05** | **1** | **-42.0372** | **-5.75024** |
| Rank 4 Rank 2 | 13.07385 | 8.37416 | 1.56121 | 0.12242 | 0.05 | 0 | -3.59125 | 29.73895 |
| Rank 4 Rank 3 | 12.22867 | 8.80283 | 1.38918 | 0.16863 | 0.05 | 0 | -5.28951 | 29.74686 |
| **Rank 5 Rank 1** | **-37.3484** | **10.83084** | **-3.44834** | **9.02E-04** | **0.05** | **1** | **-58.9025** | **-15.7944** |
| Rank 5 Rank 2 | -0.38085 | 10.21338 | -0.03729 | 0.97035 | 0.05 | 0 | -20.7061 | 19.94442 |
| Rank 5 Rank 3 | -1.22603 | 10.56771 | -0.11602 | 0.90793 | 0.05 | 0 | -22.2564 | 19.80437 |
| Rank 5 Rank 4 | -13.4547 | 11.19603 | -1.20174 | 0.23301 | 0.05 | 0 | -35.7355 | 8.82611 |

**1.4 Means comparisons of mean % dig-time in forward/positive Ranks in OPA (5-Position) Paradigm** **PT1 + RANKS**
| **Fisher LSD Test** | **MeanDiff** | **SEM** | **t Value** | **Prob** | **Alpha** | **Sig** | **LCL** | **UCL** |
| --- | --- | --- | --- | --- | --- | --- | --- | --- |
| Rank2+ Rank1+ | -17.7178 | 9.49361 | -1.86629 | 0.06697 | 0.05 | 0 | -36.7145 | 1.27888 |
| Rank3+ Rank1+ | -8.08931 | 10.26909 | -0.78773 | 0.43401 | 0.05 | 0 | -28.6377 | 12.45909 |
| Rank3+ Rank2+ | 9.62849 | 10.72107 | 0.89809 | 0.37279 | 0.05 | 0 | -11.8243 | 31.08131 |
| Rank4+ Rank1+ | -11.1686 | 12.08977 | -0.92381 | 0.35935 | 0.05 | 0 | -35.3602 | 13.02299 |
| Rank4+ Rank2+ | 6.5492 | 12.47597 | 0.52495 | 0.60159 | 0.05 | 0 | -18.4152 | 31.51356 |
| Rank4+ Rank3+ | -3.07929 | 13.07576 | -0.2355 | 0.81464 | 0.05 | 0 | -29.2438 | 23.08524 |
| **Rank5+ Rank1+** | **29.64924** | **14.47991** | **2.04761** | **0.04506** | **0.05** | **1** | **0.675** | **58.62347** |
| **Rank5+ Rank2+** | **47.36703** | **14.80388** | **3.19964** | **0.00222** | **0.05** | **1** | **17.74453** | **76.98954** |
| **Rank5+ Rank3+** | **37.73855** | **15.31276** | **2.46452** | **0.01665** | **0.05** | **1** | **7.09779** | **68.3793** |
| **Rank5+ Rank4+** | **40.81783** | **16.58882** | **2.46056** | **0.01682** | **0.05** | **1** | **7.62368** | **74.01198** |

PT3+ RANKS
| Fisher LSD Test | MeanDiff | SEM | t Value | Prob | Alpha | Sig | LCL | UCL |
| --- | --- | --- | --- | --- | --- | --- | --- | --- |
| Rank2+ Rank1+ | -6.18147 | 5.74423 | -1.07612 | 2.85E-01 | 0.05 | 0 | -17.6026 | 5.2396 |
| **Rank3+ Rank1+** | **-27.1742** | **6.33242** | **-4.29128** | **4.68E-05** | **0.05** | **1** | **-39.7648** | **-14.5837** |
| **Rank3+ Rank2+** | **-20.9927** | **6.60567** | **-3.17799** | **2.07E-03** | **0.05** | **1** | **-34.1266** | **-7.85889** |
| **Rank4+ Rank1+** | **-19.6977** | **7.49957** | **-2.62651** | **1.02E-02** | **0.05** | **1** | **-34.6088** | **-4.78653** |
| Rank4+ Rank2+ | -13.5162 | 7.73168 | -1.74816 | 0.08405 | 0.05 | 0 | -28.8889 | 1.85644 |
| Rank4+ Rank3+ | 7.47652 | 8.17815 | 0.91421 | 0.36319 | 0.05 | 0 | -8.78383 | 23.73688 |
| Rank5+ Rank1+ | -19.5173 | 10.2985 | -1.89516 | 6.15E-02 | 0.05 | 0 | -39.9935 | 0.9589 |
| Rank5+ Rank2+ | -13.3358 | 10.46874 | -1.27387 | 0.20618 | 0.05 | 0 | -34.1505 | 7.47885 |
| Rank5+ Rank3+ | 7.65692 | 10.80268 | 0.7088 | 0.48039 | 0.05 | 0 | -13.8217 | 29.13554 |
| Rank5+ Rank4+ | 0.1804 | 11.5257 | 0.01565 | 0.98755 | 0.05 | 0 | -22.7358 | 23.09658 |

PT3- RANKS
| Fisher LSD Test | MeanDiff | SEM | t Value | Prob | Alpha | Sig | LCL | UCL |
| --- | --- | --- | --- | --- | --- | --- | --- | --- |
| **Rank3- Rank2-** | **-8.54817** | **3.47055** | **-2.46306** | **0.01661** | **0.05** | **1** | **-15.488** | **-1.60838** |
| Rank4- Rank2- | -6.09446 | 3.96365 | -1.53759 | 0.12932 | 0.05 | 0 | -14.0203 | 1.83134 |
| Rank4- Rank3- | 2.45371 | 4.15711 | 0.59024 | 0.55721 | 0.05 | 0 | -5.85894 | 10.76636 |
| **Rank5- Rank2-** | **-15.074** | **5.28486** | **-2.85229** | **0.00592** | **0.05** | **1** | **-25.6417** | **-4.50623** |
| Rank5- Rank3- | -6.52579 | 5.43146 | -1.20148 | 0.23421 | 0.05 | 0 | -17.3867 | 4.33509 |
| Rank5- Rank4- | -8.9795 | 5.75904 | -1.5592 | 0.12412 | 0.05 | 0 | -20.4954 | 2.53642 |

RPT2 + RANKS
| Fisher LSD Test | MeanDiff | SEM | t Value | Prob | Alpha | Sig | LCL | UCL |
| --- | --- | --- | --- | --- | --- | --- | --- | --- |
| **Rank2+ Rank1+** | **-26.4068** | **7.40919** | **-3.56406** | **5.93E-04** | **0.05** | **1** | **-41.131** | **-11.6826** |
| **Rank3+ Rank1+** | **-30.7049** | **8.1781** | **-3.75453** | **3.11E-04** | **0.05** | **1** | **-46.9572** | **-14.4527** |
| Rank3+ Rank2+ | -4.2981 | 8.58038 | -0.50092 | 0.61768 | 0.05 | 0 | -21.3498 | 12.7536 |
| **Rank4+ Rank1+** | **-37.281** | **9.39592** | **-3.96778** | **1.48E-04** | **0.05** | **1** | **-55.9534** | **-18.6085** |
| Rank4+ Rank2+ | -10.8741 | 9.74807 | -1.11552 | 0.26767 | 0.05 | 0 | -30.2464 | 8.49812 |
| Rank4+ Rank3+ | -6.57603 | 10.34457 | -0.6357 | 0.52662 | 0.05 | 0 | -27.1337 | 13.98162 |
| **Rank5+ Rank1+** | **-36.5032** | **12.34866** | **-2.95604** | **0.004** | **0.05** | **1** | **-61.0435** | **-11.9628** |
| Rank5+ Rank2+ | -10.0964 | 12.61867 | -0.80011 | 0.4258 | 0.05 | 0 | -35.1733 | 14.98061 |
| Rank5+ Rank3+ | -5.79825 | 13.08496 | -0.44312 | 0.65876 | 0.05 | 0 | -31.8019 | 20.20535 |
| Rank5+ Rank4+ | 0.77778 | 13.87869 | 0.05604 | 0.95544 | 0.05 | 0 | -26.8032 | 28.35876 |

RPT3 + RANKS
| Fisher LSD Test | MeanDiff | SEM | t Value | Prob | Alpha | Sig | LCL | UCL |
| --- | --- | --- | --- | --- | --- | --- | --- | --- |
| **Rank2+ Rank1+** | **-46.7475** | **8.35591** | **-5.59455** | **9.27E-07** | **0.05** | **1** | **-63.5308** | **-29.9642** |
| **Rank3+ Rank1+** | **-40.6656** | **8.959** | **-4.53907** | **3.57E-05** | **0.05** | **1** | **-58.6602** | **-22.6709** |
| Rank3+ Rank2+ | 6.08195 | 9.32848 | 0.65198 | 0.5174 | 0.05 | 0 | -12.6549 | 24.81876 |
| **Rank4+ Rank1+** | **-26.0283** | **9.92663** | **-2.62207** | **0.01155** | **0.05** | **1** | **-45.9665** | **-6.09009** |
| **Rank4+ Rank2+** | **20.71919** | **10.26133** | **2.01915** | **0.04885** | **0.05** | **1** | **0.10871** | **41.32967** |
| Rank4+ Rank3+ | 14.63724 | 10.75813 | 1.36057 | 0.17975 | 0.05 | 0 | -6.9711 | 36.24557 |
| **Rank5+ Rank1+** | **-34.4555** | **11.77886** | **-2.9252** | **5.16E-03** | **0.05** | **1** | **-58.114** | **-10.797** |
| Rank5+ Rank2+ | 12.29199 | 12.06227 | 1.01904 | 0.31309 | 0.05 | 0 | -11.9358 | 36.51977 |
| Rank5+ Rank3+ | 6.21004 | 12.48763 | 0.4973 | 0.62116 | 0.05 | 0 | -18.8721 | 31.29217 |
| Rank5+ Rank4+ | -8.4272 | 13.19906 | -0.63847 | 0.52608 | 0.05 | 0 | -34.9383 | 18.08389 |

1.5 Comparison of mean % dig-time in Ranks in OPA (5-Position) Paradigm
| OFPA (5-position) Rank Means |  |  |  |  |  |  |  |  |  |
| --- | --- | --- | --- | --- | --- | --- | --- | --- | --- |
|  |  | Mean | S.E.M |  | Mean | S.E.M |  | Mean | S.E.M |
| PT1W | Rank 1 | 16.12541 | 2.82763 | Rank 1 | 16.12541 | 2.82763 |  |  |  |
|  | Rank 2 | 22.25064 | 2.71955 | Rank2+ | 21.39994 | 3.66348 | Rank2- | 23.08285 | 4.04891 |
|  | Rank 3 | 23.07762 | 3.1705 | Rank3+ | 24.96947 | 4.98712 | Rank3- | 20.89913 | 3.73065 |
|  | Rank 4 | 15.67076 | 2.80135 | Rank4+ | 17.00563 | 4.14121 | Rank4- | 14.00216 | 3.68074 |
|  | Rank 5 | 19.8489 | 4.55145 | Rank5+ | 15.28681 | 5.18422 | Rank5- | 24.06005 | 7.33864 |
| PT2W | Rank 1 | 22.36516 | 3.08733 | Rank 1 | 22.36516 | 3.08733 |  |  |  |
|  | Rank 2 | 20.35107 | 2.896 | Rank2+ | 28.03372 | 4.50661 | Rank2- | 14.33857 | 3.34032 |
|  | Rank 3 | 19.05334 | 2.62804 | Rank3+ | 18.26991 | 4.93839 | Rank3- | 19.60635 | 2.95582 |
|  | Rank 4 | 17.77082 | 4.40246 | Rank4+ | 17.77082 | 4.40246 | Rank4- | 14.76133 | 3.11803 |
|  | Rank 5 | 19.31131 | 12.2330 | Rank5+ | -- | -- | Rank5- | 19.31131 | 12.2330 |
| PT3W | Rank 1 | 43.92557 | 3.33856 | Rank 1 | 43.92557 | 3.33856 |  |  |  |
|  | Rank 2 | 14.27013 | 1.89402 | Rank2+ | 18.35628 | 3.23607 | Rank2- | 10.26911 | 1.86428 |
|  | Rank 3 | 13.52412 | 1.75571 | Rank3+ | 15.67686 | 2.76286 | Rank3- | 11.43117 | 2.16693 |
|  | Rank 4 | 15.9178 | 2.49431 | Rank4+ | 18.15293 | 4.43217 | Rank4- | 13.7758 | 2.43989 |
|  | Rank 5 | 10.62523 | 3.39137 | Rank5+ | 10.30173 | 6.53562 | Rank5- | 10.92177 | 2.91145 |
| PT10 | Rank 1 | 19.81247 | 2.71365 | Rank 1 | 19.81247 | 2.71365 |  |  |  |
|  | Rank 2 | 16.93474 | 2.00799 | Rank2+ | 12.25009 | 2.76324 | Rank2- | 21.6194 | 2.78857 |
|  | Rank 3 | 26.39512 | 3.38464 | Rank3+ | 25.04208 | 4.96128 | Rank3- | 27.67878 | 4.67426 |
|  | Rank 4 | 13.11077 | 2.56313 | Rank4+ | 11.13072 | 4.76733 | Rank4- | 14.9385 | 2.27669 |
|  | Rank 5 | 27.62096 | 6.87448 | Rank5+ | 33.45894 | 12.1016 | Rank5- | 21.78297 | 6.70667 |
| PT30 | Rank 1 | 32.07597 | 3.0774 | Rank 1 | 32.07597 | 3.0774 |  |  |  |
|  | Rank 2 | 21.06613 | 1.91341 | Rank2+ | 26.70862 | 3.46661 | Rank2- | 16.83427 | 1.92704 |
|  | Rank 3 | 13.8956 | 2.33904 | Rank3+ | 18.70939 | 3.70951 | Rank3- | 10.6864 | 2.9338 |
|  | Rank 4 | 13.31831 | 1.95763 | Rank4+ | 10.77695 | 3.72312 | Rank4- | 14.58898 | 2.28524 |
|  | Rank 5 | 14.80029 | 3.92161 | Rank5+ | -- | -- | Rank5- | 14.80029 | 3.92161 |

| OFPA 5-position Rank Analysis | 1 way-ANOVA RESULTS |  |  |  |  |
| --- | --- | --- | --- | --- | --- |
|  |  | F Value | Prob>F | DF | N |
| PT1W | Total Ranks | 1.23038 | 0.29807 | 4 | 14 |
|  | Positive Ranks | 0.98848 | 0.4153 | 4 | 14 |
|  | Negative Ranks | 0.75292 | 0.52299 | 3 | 14 |
| PT2W | Total Ranks | 0.20894 | 0.93299 | 4 | 6 |
|  | Positive Ranks | 0.73314 | 0.5367 | 3 | 6 |
|  | Negative Ranks | 0.49368 | 0.68823 | 3 | 6 |
| PT3W | **Total Ranks** | **29.53137** | **1.31E-20** | **4** | **12** |
|  | **Positive Ranks** | **14.86651** | **1.95E-10** | **4** | **12** |
|  | Negative Ranks | 0.42495 | 0.73548 | 3 | 12 |
| PT10 | **Total Ranks** | **3.41583** | **0.00942** | **4** | **14** |
|  | **Positive Ranks** | **3.09215** | **0.01712** | **4** | **14** |
|  | Negative Ranks | 1.72022 | 0.16629 | 3 | 14 |
| PT30 | **Total Ranks** | **9.0497** | **8.16E-07** | **4** | **12** |
|  | **Positive Ranks** | **4.9103** | **3.01E-03** | **3** | **12** |
|  | Negative Ranks | 1.25476 | 0.29329 | 3 | 12 |

1.6 Means comparisons of mean % dig-time in Ranks in OFPA (5-Position) Paradigm PT3W RANKS
| Fisher LSD Test | MeanDiff | SEM | t Value | Prob | Alpha | Sig | LCL | UCL |
| --- | --- | --- | --- | --- | --- | --- | --- | --- |
| **Rank 2 Rank 1** | **-29.6555** | **3.15055** | **-9.41279** | **1.56E-18** | **0.05** | **1** | **-35.8563** | **-23.4546** |
| **Rank 3 Rank 1** | **-30.4015** | **3.34834** | **-9.07956** | **1.73E-17** | **0.05** | **1** | **-36.9916** | **-23.8113** |
| Rank 3 Rank 2 | -0.74601 | 2.98179 | -0.25019 | 0.80262 | 0.05 | 0 | -6.61469 | 5.12267 |
| **Rank 4 Rank 1** | **-28.0078** | **3.71613** | **-7.53681** | **6.19E-13** | **0.05** | **1** | **-35.3218** | **-20.6938** |
| Rank 4 Rank 2 | 1.64767 | 3.38959 | 0.4861 | 0.62727 | 0.05 | 0 | -5.02364 | 8.31898 |
| Rank 4 Rank 3 | 2.39368 | 3.57418 | 0.66971 | 0.50357 | 0.05 | 0 | -4.64094 | 9.4283 |
| **Rank 5 Rank 1** | **-33.3004** | **4.6723** | **-7.12719** | **8.16E-12** | **0.05** | **1** | **-42.4963** | **-24.1044** |
| Rank 5 Rank 2 | -3.6449 | 4.41701 | -0.8252 | 0.40994 | 0.05 | 0 | -12.3384 | 5.04857 |
| Rank 5 Rank 3 | -2.89889 | 4.5602 | -0.63569 | 0.52548 | 0.05 | 0 | -11.8742 | 6.0764 |
| Rank 5 Rank 4 | -5.29257 | 4.8367 | -1.09425 | 0.27475 | 0.05 | 0 | -14.8121 | 4.22692 |

PT10 RANKS
| Fisher LSD Test | MeanDiff | SEM | t Value | Prob | Alpha | Sig | LCL | UCL |
| --- | --- | --- | --- | --- | --- | --- | --- | --- |
| Rank 2 Rank 1 | -2.87772 | 3.83738 | -0.74992 | 0.45387 | 0.05 | 0 | -10.4283 | 4.67288 |
| Rank 3 Rank 1 | 6.58266 | 4.08031 | 1.61327 | 0.1077 | 0.05 | 0 | -1.44595 | 14.61126 |
| **Rank 3 Rank 2** | **9.46038** | **3.62882** | **2.60701** | **0.00958** | **0.05** | **1** | **2.32014** | **16.60062** |
| Rank 4 Rank 1 | -6.7017 | 4.53572 | -1.47754 | 0.14055 | 0.05 | 0 | -15.6264 | 2.223 |
| Rank 4 Rank 2 | -3.82398 | 4.13428 | -0.92494 | 0.35571 | 0.05 | 0 | -11.9588 | 4.31081 |
| **Rank 4 Rank 3** | **-13.2844** | **4.3607** | **-3.04638** | **0.00252** | **0.05** | **1** | **-21.8647** | **-4.70404** |
| Rank 5 Rank 1 | 7.80849 | 5.74442 | 1.35932 | 0.17503 | 0.05 | 0 | -3.4945 | 19.11148 |
| Rank 5 Rank 2 | 10.68621 | 5.43303 | 1.9669 | 0.05009 | 0.05 | 0 | -0.00407 | 21.3765 |
| Rank 5 Rank 3 | 1.22583 | 5.60726 | 0.21862 | 0.82709 | 0.05 | 0 | -9.80726 | 12.25892 |
| **Rank 5 Rank 4** | **14.51019** | **5.94686** | **2.43997** | **0.01525** | **0.05** | **1** | **2.80887** | **26.21151** |

PT30 RANKS
| Fisher LSD Test | MeanDiff | SEM | t Value | Prob | Alpha | Sig | LCL | UCL |
| --- | --- | --- | --- | --- | --- | --- | --- | --- |
| **Rank 2 Rank 1** | **-11.0098** | **3.19155** | **-3.44968** | **6.65E-04** | **0.05** | **1** | **-17.2975** | **-4.72212** |
| **Rank 3 Rank 1** | **-18.1804** | **3.41579** | **-5.32245** | **2.39E-07** | **0.05** | **1** | **-24.9099** | **-11.4509** |
| **Rank 3 Rank 2** | **-7.17054** | **2.98154** | **-2.40497** | **0.01695** | **0.05** | **1** | **-13.0445** | **-1.29657** |
| **Rank 4 Rank 1** | **-18.7577** | **3.88905** | **-4.8232** | **2.54E-06** | **0.05** | **1** | **-26.4195** | **-11.0958** |
| **Rank 4 Rank 2** | **-7.74783** | **3.51378** | **-2.20498** | **0.02842** | **0.05** | **1** | **-14.6704** | **-0.82529** |
| Rank 4 Rank 3 | -0.57729 | 3.71864 | -0.15524 | 0.87676 | 0.05 | 0 | -7.90342 | 6.74884 |
| **Rank 5 Rank 1** | **-17.2757** | **5.69298** | **-3.03456** | **0.00268** | **0.05** | **1** | **-28.4915** | **-6.05988** |
| Rank 5 Rank 2 | -6.26584 | 5.44353 | -1.15106 | 0.25088 | 0.05 | 0 | -16.9902 | 4.45851 |
| Rank 5 Rank 3 | 0.9047 | 5.57796 | 0.16219 | 0.87129 | 0.05 | 0 | -10.0845 | 11.89389 |
| Rank 5 Rank 4 | 1.48199 | 5.87968 | 0.25205 | 0.80122 | 0.05 | 0 | -10.1016 | 13.06562 |

1.7 Means comparisons of mean % dig-time in forward/positive Ranks in OFPA (5-Position) Paradigm PT3W + RANKS
| Fisher LSD Test | MeanDiff | SEM | t Value | Prob | Alpha | Sig | LCL | UCL |
| --- | --- | --- | --- | --- | --- | --- | --- | --- |
| Rank 2+ Rank 1 | **-25.5693** | **4.35907** | **-5.86577** | **2.28E-08** | **0.05** | **1** | **-34.1742** | **-16.9644** |
| Rank 3+ Rank 1 | **-28.2487** | **4.75685** | **-5.93853** | **1.58E-08** | **0.05** | **1** | **-37.6388** | **-18.8586** |
| Rank 3+ Rank 2+ | -2.67941 | 4.97782 | -0.53827 | 0.59109 | 0.05 | 0 | -12.5057 | 7.1469 |
| Rank 4+ Rank 1 | **-25.7726** | **5.48066** | **-4.70247** | **5.29E-06** | **0.05** | **1** | **-36.5916** | **-14.9537** |
| Rank 4+ Rank 2+ | -0.20335 | 5.67351 | -0.03584 | 0.97145 | 0.05 | 0 | -11.403 | 10.99626 |
| Rank 4+ Rank 3+ | 2.47607 | 5.98455 | 0.41374 | 0.67958 | 0.05 | 0 | -9.33754 | 14.28967 |
| Rank 5+ Rank 1 | **-33.6238** | **7.32221** | **-4.59203** | **8.51E-06** | **0.05** | **1** | **-48.078** | **-19.1697** |
| Rank 5+ Rank 2+ | -8.05455 | 7.46766 | -1.07859 | 0.2823 | 0.05 | 0 | -22.7958 | 6.68673 |
| Rank 5+ Rank 3+ | -5.37513 | 7.70662 | -0.69747 | 0.48646 | 0.05 | 0 | -20.5881 | 9.83787 |
| Rank 5+ Rank 4+ | -7.8512 | 8.17325 | -0.9606 | 0.33812 | 0.05 | 0 | -23.9853 | 8.28293 |

PT10 + RANKS
| Fisher LSD Test | MeanDiff | SEM | t Value | Prob | Alpha | Sig | LCL | UCL |
| --- | --- | --- | --- | --- | --- | --- | --- | --- |
| Rank 2+ Rank 1 | -7.56237 | 4.6914 | -1.61196 | 0.1087 | 0.05 | 0 | -16.8189 | 1.69415 |
| Rank 3+ Rank 1 | 5.22961 | 5.15863 | 1.01376 | 0.31204 | 0.05 | 0 | -4.9488 | 15.40802 |
| Rank 3+ Rank 2+ | **12.79199** | **5.3785** | **2.37836** | **0.01843** | **0.05** | **1** | **2.17976** | **23.40421** |
| Rank 4+ Rank 1 | -8.68174 | 5.97433 | -1.45317 | 0.1479 | 0.05 | 0 | -20.4696 | 3.10611 |
| Rank 4+ Rank 2+ | -1.11937 | 6.16517 | -0.18156 | 0.85613 | 0.05 | 0 | -13.2838 | 11.04504 |
| Rank 4+ Rank 3+ | **-13.9114** | **6.52776** | **-2.13111** | **0.03442** | **0.05** | **1** | **-26.7912** | **-1.03155** |
| Rank 5+ Rank 1 | 13.64648 | 7.84468 | 1.73958 | 0.08362 | 0.05 | 0 | -1.83174 | 29.12469 |
| Rank 5+ Rank 2+ | **21.20885** | **7.99098** | **2.6541** | **0.00866** | **0.05** | **1** | **5.44197** | **36.97572** |
| Rank 5+ Rank 3+ | 8.41686 | 8.27393 | 1.01727 | 0.31037 | 0.05 | 0 | -7.9083 | 24.74203 |
| Rank 5+ Rank 4+ | **22.32822** | **8.80563** | **2.53568** | **0.01206** | **0.05** | **1** | **4.95397** | **39.70247** |

PT30 + RANKS
| Fisher LSD Test | MeanDiff | SEM | t Value | Prob | Alpha | Sig | LCL | UCL |
| --- | --- | --- | --- | --- | --- | --- | --- | --- |
| Rank 2+ Rank 1 | -5.36735 | 4.38732 | -1.22338 | 2.24E-01 | 0.05 | 0 | -14.057 | 3.3223 |
| Rank 3+ Rank 1 | **-13.3666** | **4.97476** | **-2.68688** | **8.27E-03** | **0.05** | **1** | **-23.2197** | **-3.51345** |
| Rank 3+ Rank 2+ | -7.99923 | 5.24385 | -1.52545 | 0.12987 | 0.05 | 0 | -18.3853 | 2.38688 |
| Rank 4+ Rank 1 | **-21.299** | **6.42238** | **-3.31637** | **1.22E-03** | **0.05** | **1** | **-34.0194** | **-8.57868** |
| Rank 4+ Rank 2+ | **-15.9317** | **6.63301** | **-2.40188** | **0.0179** | **0.05** | **1** | **-29.0692** | **-2.79416** |
| Rank 4+ Rank 3+ | -7.93244 | 7.03537 | -1.12751 | 0.26185 | 0.05 | 0 | -21.8669 | 6.00199 |

## Notes

### Competing Interest Statement

The authors have declared no competing interest.

### Summary of Updates

Author names were miss spelt and now has been corrected.

